# Single nucleus multi-omic atlas of human dorsal root ganglia reveals the contribution of non-neuronal cell types to pain

**DOI:** 10.1101/2025.09.21.677655

**Authors:** Kevin Boyer, Pauline Meriau, Lite Yang, Huma Naz, Sarah Rosen, George Murray, Adam J. Dourson, Juliet M. Mwirigi, John Del Rosario, Maria Payne, Prashant Gupta, Jiwon Yi, Richard A. Slivicki, Zachariah Bertels, John Lemen, Yujie Chen, Ting Wang, Alexander Chamessian, Bryan A. Copits, Robert W. Gereau, Valeria Cavalli, Guoyan Zhao

## Abstract

Sensory neurons residing in dorsal root ganglia (DRG) transmit sensory information such as pain, itch, touch, pressure and bodily position to the central nervous system. The activity of sensory neurons is regulated by non-neuronal cell types in the DRG, including satellite glial cells (SGCs), immune cells and fibroblasts. Dysregulated gene expression in DRG cells contributes to sensory nervous system disorders such as chronic pain. Understanding the genetic underpinnings of these conditions requires dissecting transcriptional regulation in human tissue. In this study, we profiled transcriptomic and chromatin accessibility landscapes from postmortem human DRG (hDRG) samples at single-nucleus level. We demonstrate that sequencing depth significantly impacts downstream analysis, with deeper sequencing yielding more detected cells and features, improved data integration, refined clustering and annotation, and more accurate scientific interpretations. We identified nine major cell types, defined their molecular signatures, and mapped cis-regulatory landscapes. Integration of gene expression with chromatin accessibility enabled peak-gene association and transcriptional network analyses, revealing transcription factors, their target genes, regulatory elements and potential partners that cooperatively drive cell-type-specific gene expression programs. This integrative approach identified cell types, genes, and cis-regulatory regions potentially driving pain conditions. Our unbiased genome-wide analysis not only recovered known pain-related genes but also highlighted novel candidate genes and regulatory regions implicated in pain mechanisms. Importantly, our results demonstrate that non-neuronal cells, including endothelial cells, fibroblasts, macrophages, and SGCs, play critical roles in pain pathogenesis and should be investigated as therapeutic targets.

**One Sentence Summary:** Our work provides a comprehensive single-nucleus multi-omic atlas of human dorsal root ganglia, uncovering novel cell-type-specific regulatory mechanisms and candidate therapeutic targets for pain, thereby directly advancing translational insights into human sensory disorders and chronic pain pathogenesis.

## INTRODUCTION

Vertebrates sense a wide range of external and internal stimuli through functionally distinct peripheral sensory neurons whose cell bodies reside in DRG. These neurons are tuned to respond to specific physical and chemical stimuli, enabling the perception and discrimination of various sensations, such as touch, itch, pain, temperature, and proprioception(*1*). Transmission of sensory information requires the integrated function of multiple cell types including SGCs, fibroblasts, macrophages, and Schwann cells. In rodent models, cellular and gene expression changes in these cells have been implicated in various painful states(*2–8*) and degenerative diseases(*9–11*). The last three decades have seen significant advances in understanding the electrochemical, cellular and molecular characteristics of neurons and non-neuronal cells in DRG of animal models(*7, 8, 12–14*). Recent advances in single-cell or single-nucleus RNA sequencing technology (sc/snRNA-seq) have greatly expanded our knowledge of DRG cell type composition and gene expression in rodent models(*15–19*). Recent studies have revealed important similarities and differences in molecular and cellular characteristics of DRG between animal models and human(*17, 20–25*), which have profound implications for translating data from rodent models to human pathologies, and subsequent therapeutic developments. However, how the identity of different cell types in human (hDRG) are regulated and how they may contribute to the development of pain remains poorly understood.

Chronic pain, conventionally defined as pain lasting longer than 3 months, is highly prevalent worldwide and represents a significant socioeconomic and public health burden(*26, 27*), contributing to excess mortality(*28, 29*). At present, the mechanisms underlying chronic pain in various conditions are not fully understood, and existing treatments often prove ineffective for most patients(*30–34*). The widespread use of opioids, the most commonly prescribed class of pain medication, has sparked a nationwide crisis. Therefore, a better understanding of the cellular and molecular mechanisms underlying pain conditions may pave the way for the development of novel and safer treatments for many chronic pain conditions. Twin studies have indicated that chronic pain conditions show heritability ranges from 16% to 50%(*35*). Multiple large-scale genome-wide association studies (GWAS) have revealed that chronic pain conditions are heritable, with an estimated genetic contribution accounting to 10.2 to 45% heritability(*26, 34*). Furthermore, shared heritability and genetic variants across multiple chronic pain conditions have been reported (*34, 36–38*), implicating a possible shared genetic basis of pain conditions. GWAS studies have identified tens to thousands of single nucleotide polymorphisms (SNPs) associated with pain conditions. However, these studies have not directly identified causal variants and/or the genes driving chronic pain conditions. Furthermore, given that over 90% of disease- and trait-associated SNPs are located in the non-coding regions of the genome(*39, 40*), the cell type and the mechanisms through which they affect gene expression in the context of pain conditions have not yet been described.

In this study, we analyzed 11 human DRG samples from seven donors (**Table S1**), using the 10x Genomics Single Cell Multiome ATAC + Gene Expression Assay, which integrates snRNA-seq and single nucleus assay for transposase-accessible chromatin sequencing (snATAC-seq) data from the same cells (**Fig. 1A**). The joint profiling of gene expression and chromatin accessibility of the same exact cells is superior at refining cell types and revealing gene regulatory networks, providing the most direct link between cis-regulatory elements and their target genes (*41*). We identified nine major cell types, defined their molecular signatures, and unveiled transcriptional network that regulates major cell type identities. We then integrated the transcriptomic and epigenetic data from single-cell multi-omics analysis of human DRG with pain-associated genes and SNPs from multiple GWAS to infer susceptible cell types and causal SNPs underlying human pain disorders. We identified cell-type-specific pain-associated genes and cis-regulatory regions potentially driving pain conditions including known pain-related genes and potential novel candidates. Our work provides new insights into the genetics and underlying biology of chronic pain and may help to inform new treatment strategies.

**Figure 1:**
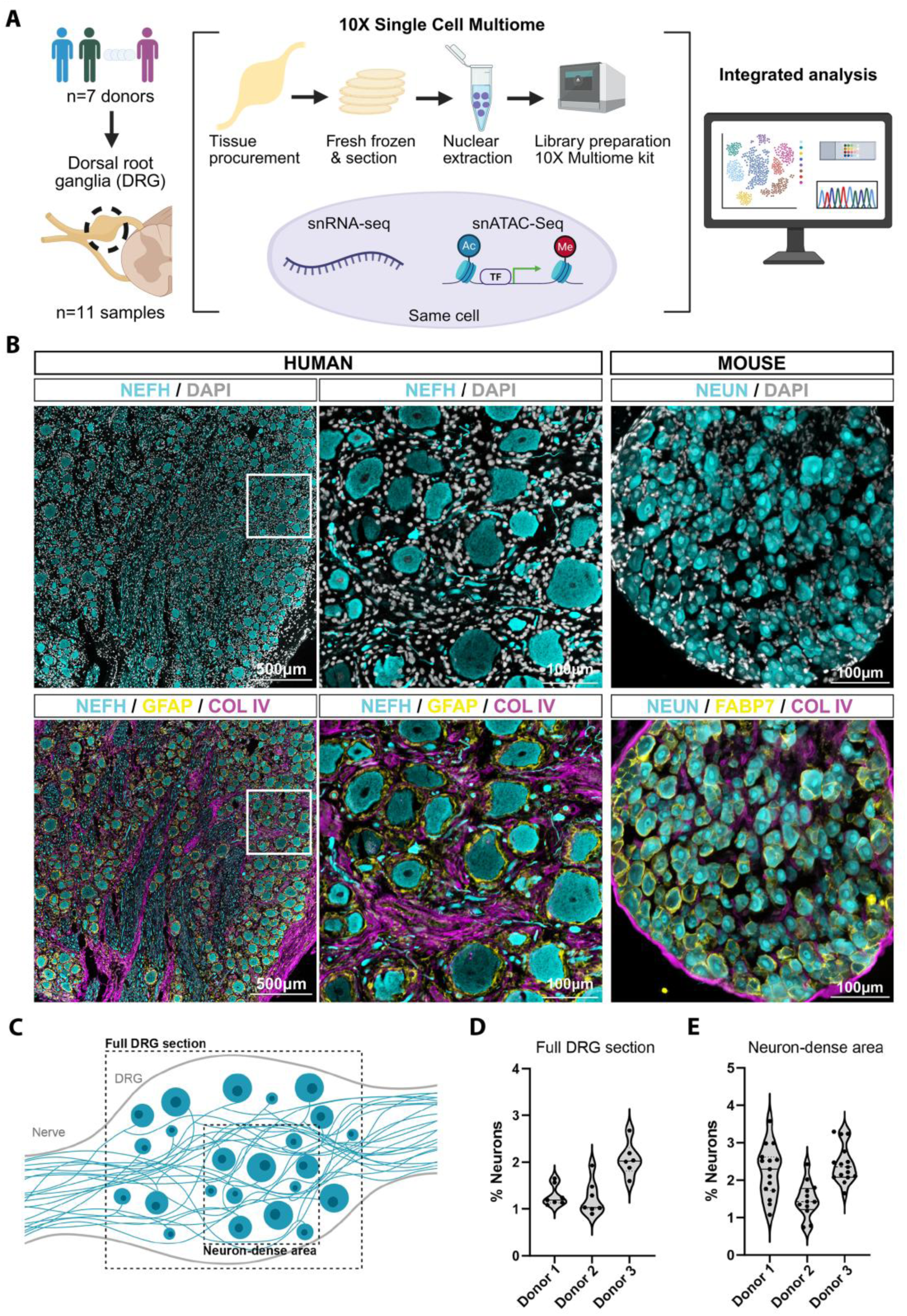
Study design and neuronal population quantification. (**A**) Workflow schematic representation illustrates the processing of hDRG, which enables the concurrent execution of snRNA-Seq and snATAC-Seq on a single cell, utilizing 10X Single Cell Multiome technology. (**B**) Representative images of immunofluorescence staining of human and mouse DRG sections. Neurons (NEFH for humans and NeuN for mice) are labeled in cyan and DAPI is shown in white. The glial cells (GFAP for humans and Fabp7 for mice) are labeled in yellow. The connective tissue (Collagen IV) is labeled in magenta. Scale bar: 500 mm (left panel) and 100 mm (middle and right panel). (**C**) A schematic representation of the distribution of neurons in an hDRG. (**D**) Quantification of the proportion of neurons in a full DRG section, including areas dense with neurons soma, and axon bundles. (**E**) Quantification of the proportion of neurons in neuron-soma-dense areas only.

## RESULTS

### Low abundance of neurons in the human DRG

The DRG comprises many different cell types including neurons, SGCs, myelinating and non-myelinating Schwann cells (mySC and nmSC), fibroblasts, endothelial cells, and immune cells. In fact, most cells in the DRG are non-neuronal, which presents a challenge for analyzing neurons in this tissue via single-cell modalities. Our previous work showed that although the actual percentage of neuronal cells in the mouse DRG is about 12%, the percentage of neurons detected in scRNA-seq and snRNA-seq analyses was only about 1% for mouse and 0.5% for rat samples (*17, 42*). In humans, DRGs also contain more neurons per DRG than animals, estimated to be 70,000 neurons in young humans versus 15,000 in rats (*43–45*). In comparison to rodents, hDRGs exhibit increased size and contain a greater number of cells (*46*). In a comparative immunofluorescence of human and mouse DRG sections, we confirmed these differences (**Fig 1B**). DRG sections stained for the neuronal markers (NEFH for human, NeuN for mouse) and for DAPI (all nuclei) revealed that the majority of nuclei in hDRG labeled non-neuronal cells, with neurons representing ∼2% of all hDRG cells (**Fig. 1C-E**), which is six times lower than the percentage observed in mice (*42*). Additionally, staining for glial cells (GFAP for human, Fabp7 for mouse) and for connective tissue (Collagen IV) demonstrated that hDRGs contain a markedly higher proportion of connective tissue compared to mouse DRGs (**Fig. 1B**), thereby highlighting key structural differences between species (*46*).

### Optimization of neuronal nuclei recovery for human DRG snRNA-seq analysis

The size of neuronal soma poses unique challenges in the unbiased characterization of their molecular profiles. As the largest cells in the DRG, neurons in humans have diameters ranging from 20 to 120µm with large nuclei(*43, 47*). Given the 10X Genomics technology, which has a maximum cell/nucleus size of 30 µm during microfluidic-based cell barcoding step, we opted for single nucleus sequencing. To determine potential limiting factors and optimize a protocol that enriches for neuronal nuclei, we first compared multiple protocols with varying extraction and treatment conditions to determine their effect on neuronal nuclei yield. We compared four nuclear dissociation methods previously described (*48*), including a nuclear extraction protocol with density gradient centrifugation alone (Opti), Opti followed by FACS (Opti/FACS), a crude extraction protocol followed by FACS (Non-gradient/FACS), and the nuclear isolation kit developed by 10X Genomics (10X kit). We also compared two different douncing conditions and two different detergents (**Table S2**). Because the proportion of neurons can differ significantly between donors, tissues sampled from different lumbar levels, and even different sections, we pooled multiple samples from multiple donors to use as starting material. This ensured that any observed differences in the proportion of cell populations are more likely to reflect the differences in cell isolation protocols. To assess neuronal nuclei enrichment, we performed snMultiome-seq on nuclei isolated by each protocol and performed preliminary unsupervised clustering and quality control using the single cell genomic tool Seurat (*49*). We identified neuronal clusters based on the unique expression of neuronal marker genes *RBFOX1* and *SNAP25* and calculated the percentage of neuronal nuclei in each sample. The nuclear extraction method using gradient centrifugation (Opti-prep) yielded the most enrichment of neuronal nuclei (2.39 ± 1.03%, n=4) with stable performance across different detergents and disassociation methods, whereas non-gradient/FACS, Opti/FACS, and 10X kit protocols yielded 0%, 0.06%, and 0.56% (n=2) neuronal nuclei, respectively (**Table S2**). Opti-prep thus preserved more neuronal nuclei compared to other nuclear extraction approaches and yielded a number of neurons comparable to the neuronal percentage measured in hDRG sections. Therefore, we used this approach to prepare nuclei for snMultiome-seq in this study.

### Impact of sequencing depth on cell annotation and data interpretation

Next, we used the optimized Opti-prep protocol on two samples to determine the impact of sequencing depth on cell type annotation and feature detection. To maximize the differences between samples, we chose two hDRG samples (Donor 1:20220418A and Donor 2:20220805A) from different lumbar levels and from two individuals with different pain histories and distinct clinical features (**Table S1**). We generated data with a target of 50k reads/cell in an initial shallow sequencing run (referred to as “shallow data”). We next performed additional sequencing with a target of total 150k reads/cell for snRNA-seq data and 100k reads/cell for snATAC-seq data, after combining data from both sequencing runs (referred to as “deep data”). We first compared the mapping statistics and sample quality assessment measurements generated by the 10X Cell Ranger ARC software (**Table S3**). We also performed *de novo* peak calling using Model-based Analysis of ChIP-Seq (MACS2)(*50*) yielding similar number of ATAC-seq peaks. As expected, increased sequencing depth resulted in more cells and more features detected in both transcripts and ATAC peaks for both samples (**Fig. S1**). The sequencing depth had a more significant impact on the median number of features detected for the snATAC-seq than that of the snRNA-seq, doubling the number of features detected for both peak-calling methods. After removing low-quality cells, we obtained 16,888 and 18,416 nuclei for shallow and deep sequencing data respectively. Our analysis thus revealed that increased sequencing depth led to a 9% increase in the number of cells.

To further compare shallow and deep sequencing data, we analyzed these two datasets independently using the same analysis pipeline and applying identical analysis parameters. First, we performed data normalization, batch effect removal, and integration for snRNA-seq and snATAC-seq data separately using Seurat v5 (*51*) and Signac v1.6.0 (*52*). Subsequently, the two single-modality datasets were projected onto a joint neighbor graph using the weighted nearest neighbor (WNN) method which enables the joint analyses of both DNA accessibility and gene expression. The shallow data resulted in 19 total clusters (**Fig. 2A**), while the deep data resulted in 18 cell clusters (**Fig. 2B**). We noticed that in shallow sequencing data, cluster 5 and 9 were largely segregated by donors with cluster 5 (referred to as C5s) cells largely from donor 1 and cluster 9 (referred to as C9s) cells from donor 2 (**Fig. 2A**). However, both clusters uniquely enriched for *PLAC8* expression (**Fig. 2C**), suggesting similar cell types shared by the two donors. We projected the cells from C5s and C9s (referred to as C5s+C9s cells) from the shallow data (**Fig. 2C**) to the UMAP of the deep sequencing data and found that most of the C5s+C9s cells (95.8% of C5s and 70.7% of C9s, mapped to a single cluster of C5 (referred to as C5d) in deep sequencing data (**Fig. 2D,E**), which is uniquely enriched for *PLAC8* expression. Additionally, C5s+C9s cells from the two donors were better aligned in deep sequencing (**Fig. 2B**), supporting that C5s and C9s from shallow data were the corresponding shared C5d cells in the two donors. Furthermore, we noticed that 40 (4.2%) cells from shallow C5s and 182 (∼29.3%) cells from shallow C9s cluster were not mapped to the deep C5d cluster. Instead, most of these cells were mapped to the C9d cluster of the deep data, which was distributed across the UMAP space and found to be a doublet cluster (**Fig. 2F**). In summary, this analysis reveals that deep sequencing enabled the detection of more cells, more features, better data integration, better cell clustering and annotation, as well as more accurate downstream analysis and scientific interpretations. Consequently, we performed deep sequencing for the high-quality samples (S1-S7) and four additional samples to a target of 150k reads/cell to assemble a high-quality single-nucleus multi-omic atlas of hDRGs (**Table S4**).

**Figure 2:**
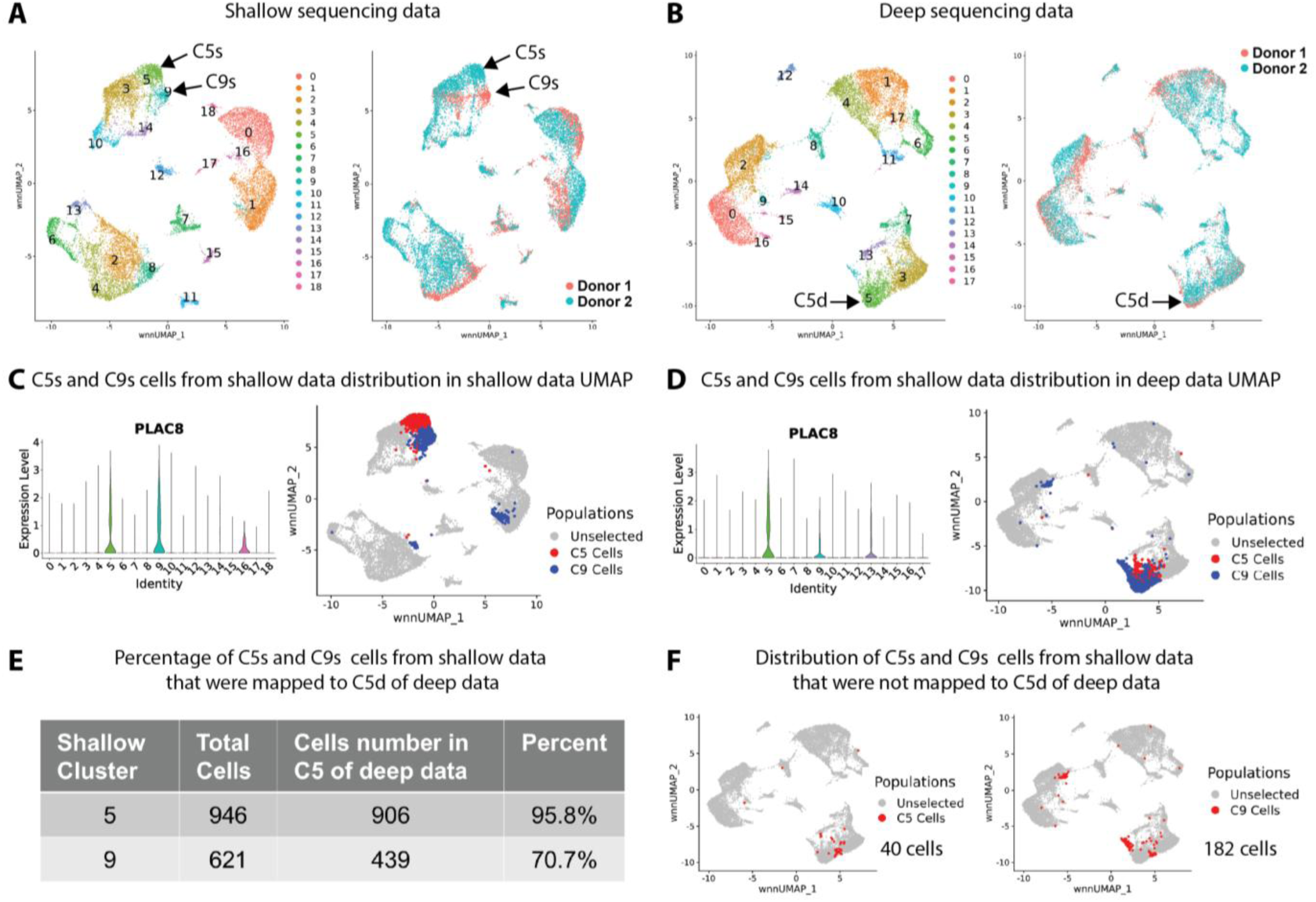
Impact of sequencing depth on cell annotation and data interpretation. (**A**) For shallow sequencing data, cells were visualized using weighted nearest neighbor (WNN) uniform manifold approximation and projection (wnnUMAP) after quality control. (left) Cells colored by cluster identity; (right) Cells colored by donor identity. (**B**) Plots for deep sequencing data similar to (a). (**C**) Violin plot of PLAC8 expression levels (left) and the distribution of cells from C5s and C9s in shallow sequencing data wnnUMAP (right). (**D**) Violin plot of PLAC8 expression levels (left) and the distribution of cells from C5s and C9s in deep sequencing data wnnUMAP (right). (**E**) A table showing the percentage of C5s and C9s cells from shallow data that were mapped to C5d of deep sequencing data. (**F**) wnnUMAP showing the distribution of C5s (left) and C9s (right) cells from shallow data that were not mapped to C5d of deep data.

### Single-nucleus multi-omic atlas of human DRGs

A total of 97,643 nuclei were obtained from 11 samples derived from 7 donors. Following the implementation of a rigorous quality control protocol, which entailed the elimination of doublets and the removal of low-quality cell clusters, a total of 66,599 high-quality transcriptome profiles were obtained. The data then underwent a series of processing steps, including data normalization, batch effect removal, integration of snRNA-seq and snATAC-seq data and projected data onto a joint neighbor graph using the WNN method as previous published(*53*) (**Fig. 3A**). We delineated 14 cell clusters with distinct gene expression profiles. By leveraging known cell type-specific marker gene expression, we identified nine major cell types including endothelial cells (referred to as Endo herein) (*FLT1, EMCN, ANO2*)(*54*), fibroblast/mesenchymal cells (Fibro) (*EGFR, APOD, DCN, PDGFRA, COL1A1, FBLN1, EYA1*), macrophages (Mac) (*SYK, OLR1, PALD1, PLAC8, F13A1, MRC1, CD163, LYVE1, SIGLEC1*), neurons (*SNAP25, RBFOX1, ROBO2, SYT1, THY1, TUBB3, PRPH*), myelinating Schwann cells (mySC) (*PRX, DRP2, NCMAP, MPZ*), non-myelinating Schwann cells (nmSC) (*NRXN1, SCN7A, SORCS3, SEMA3E*), satellite glia cells (SGCs) (*SHANK2, KCNJ3, CTNND2, FABP7*), mural cells (*PDGFRB, RGS5, MYOCD*), and T cells (*SKAP1, BCL11B, CD2, CD247*) (**Fig. 3A-B, S2a, Table S5**)(*17, 55–58*).

**Figure 3.**
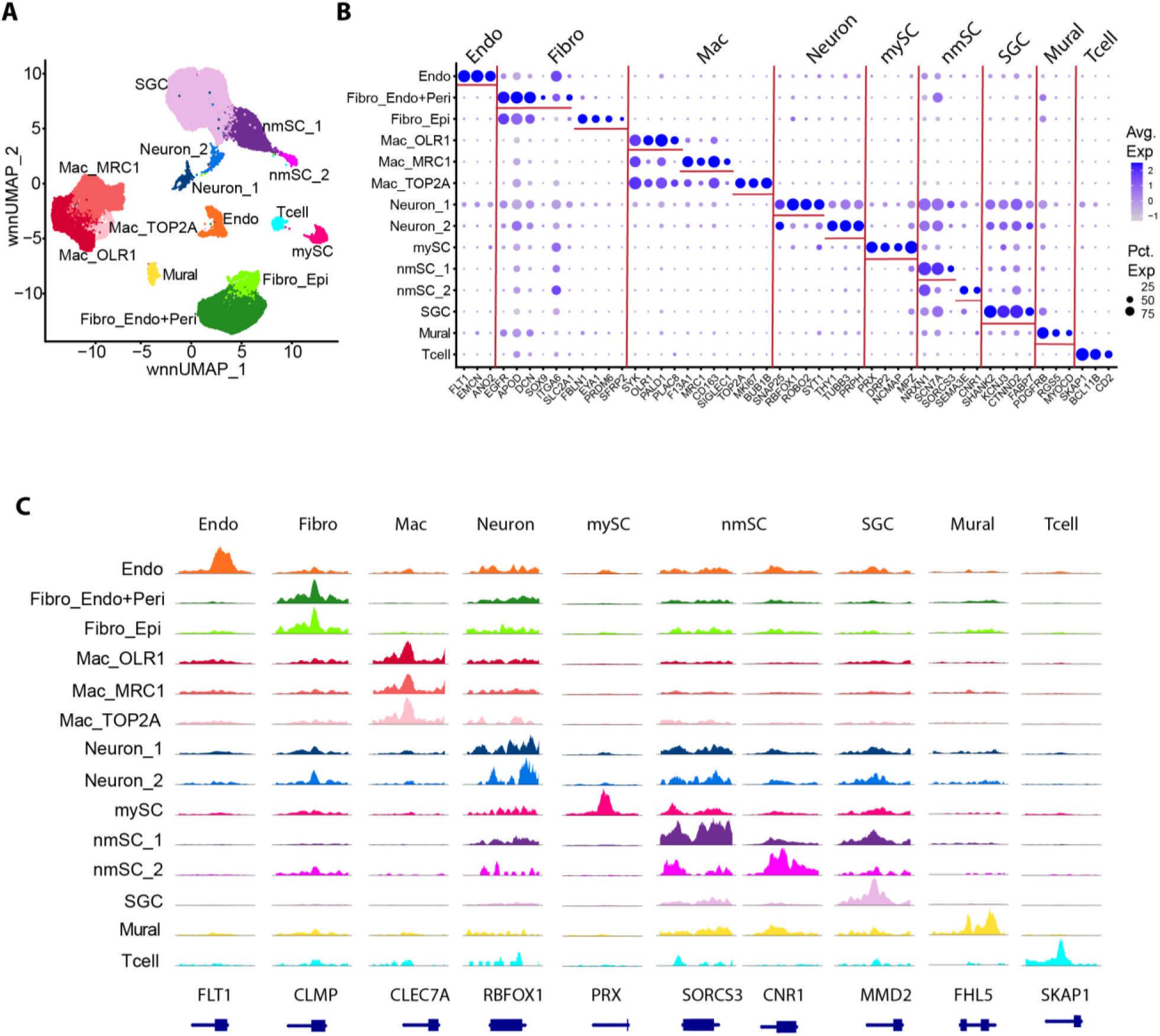
Cell type identification in human DRG of all the cells obtained from 11 samples after quality control. (**A**) Cells were visualized using wnnUMAP colored by cluster identity. (**B**) Dot plot of marker gene expression levels. (**C**) Genome browser views showing the aggregated and normalized snATAC-seq signals at promoter regions (±1 kb from transcription starts site) of selected cell-type marker genes. Tracks are colored with the same color code as in (*A)* and (*C)*.

To assess donor- and sample-based batch effects, we examined the distribution of nuclei across cell clusters for each donor, revealing an overall even distribution across cell type clusters (**Fig. S2B**). The 11 samples were distributed uniformly across cell type clusters, exhibiting equivalent numbers of unique molecular identifiers (UMIs) and detected genes (**Fig. S2C**). SGCs were the most prevalent cell type in our dataset with 21,527 nuclei, followed by macrophages with 16,441 nuclei. In total, we obtained 1,125 neurons with an average percentage of 1.7%, consistent with our antibody staining-based quantification (**Fig. 1D, E**). However, the percentage of neurons and other cell types varied significantly across samples (**Fig. S2D**), which could reflect sampling bias of neuron-sparse vs. neuron-dense areas of the DRGs or other donor variable such as age, medical history, and tissue preservation.

Human DRGs contain up to 16 neuronal subtypes (*20, 23*), and our snRNA-seq transcriptomic profile identified two neuronal populations (Neuron_1, Neuron_2), both expressing pan-neuronal markers *SNAP25*, neurofilaments (*NEFH, NEFL, NEFM*), α- and β-tubulin (*TUBA1A, TUBA1B*, *TUBB3*) (**Fig. 3B, 4A, S3A**) (*23, 46*). Both neuronal populations also expressed known peripheral sensory neuronal markers *SLC17A6* (*VGLUT2*), *SYP* (synaptophysin), and ubiquitin C-terminal hydrolase *UCHL1* (*PGP9.5*)(*23*) (**Fig. S3B**), confirming their sensory neuron identity. Mammalian DRG neurons are broadly classified into two major categories: myelinated large-diameter (A-fiber) and unmyelinated small-diameter (C-fiber) neurons(*59*). Neuron_1 population had higher expression of *RBFOX1, ROBO2,* and *SYT1* as well as unique expressions of *CACNA1B, SRRM3, NTRK1, TMEM255A, PRDM12, TRPV1,* and *TRPA1* (**Fig. 3B, 4A, S3C**). Since *Rbfox1* and *Robo2* are enriched in small diameter neurons in rat and mouse DRGs (*60, 61*), Neuron_1 cluster likely included multiple subtypes of unmyelinated small-diameter C-fibers neurons. Furthermore, Neuron_1 was enriched for *PRDM12*, a transcriptional regulator critical for the development of human C-fibers and pain-sensing neurons (nociceptors)(*62*), as well as *TRPV1* and *TRPA1,* genes known to be expressed in C-fiber nociceptors in mice required for heat and cold sensation, respectively (*1*), and *NTRK1* (TrkA), which is restricted to nociceptors (*63*). In contrast, Neuron_2 exhibited higher expression of neurofilament heavy chain *NEFH* and peripherin (*PRPH*) **(Fig. 4A)**. Given that NEFH, but not PRPH, is enriched in large-diameter A-fiber DRG neurons (*23*), Neuron_2 likely represents a mixture of small and large myelinated neurons. Given the limited number of neurons recovered in our dataset, we were underpowered to resolve the full heterogeneity of neuronal subtypes (*20, 23*).

**Figure 4:**
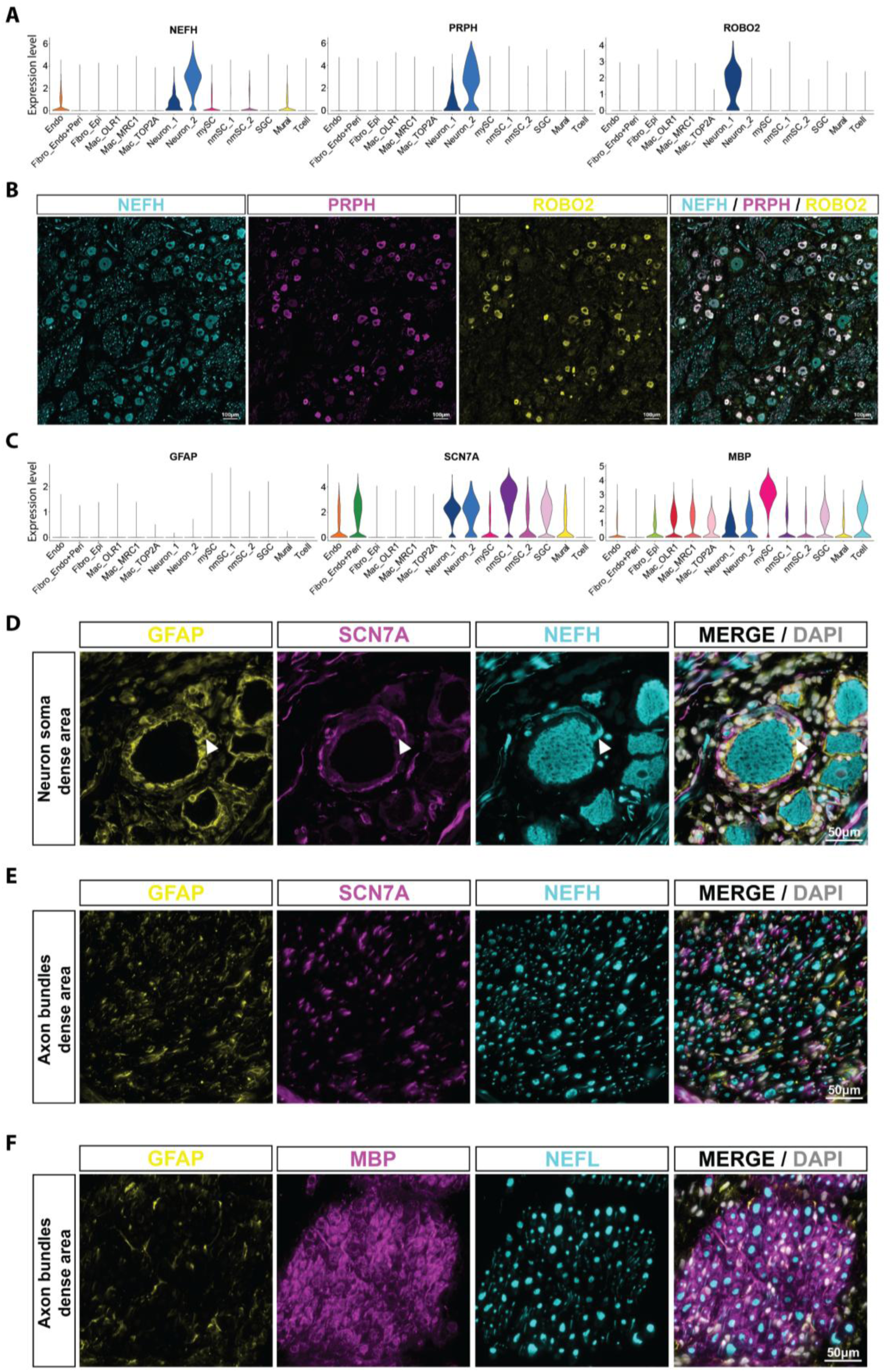
Examination of neuronal and glia cell marker protein expressions in human DRGs. (**A**) Violin-plots showing the expression of NEFH, ROBO2 and PRPH across the different cell populations. (**B**) Representative images of immunofluorescence staining of neurons, labeled for NEFH (cyan), PRPH (magenta) and ROBO2 (yellow). Scale bar: 100 mm. (**C**) Violin-plots showing the expression of GFAP, SCN7A and MBP across the different cell populations. Representative images of immunofluorescence staining of neuron soma dense area (**D**), or axon bundles dense area (**E**) of hDRG, labeled for GFAP (yellow), SCN7A (magenta) NEFH (cyan), and DAPI (gray). Scale bar: 50 mm. (**f**F Representative images of immunofluorescence staining of axon bundles dense area of hDRG, labeled for MBP (yellow), SCN7A (magenta) NEFH (cyan), and DAPI (gray).

We next examined expression of markers for the two neuronal populations at the protein level, given that correlations between mRNA and protein expression are not always consistent (*64–67*). Immunofluorescence staining with antibodies against NEFH and PRPH (transcriptionally enriched in Neuron_2) and ROBO2 (transcriptionally enriched in Neuron_1) revealed striking discrepancies between protein and transcript expression. All neurons displayed comparable NEFH protein levels, consistent with prior report (*68*), but not with *NEFH* mRNA expression profile (**Fig. 4A, B**). PRPH and ROBO2 co-localized in smaller neurons (likely C-fibers), consistent with their transcript expression profile in different neurons (**Fig. 4A, B)**. However, PRPH proteins were absent from larger neurons, in contrast to the observed higher expression at mRNA level. The difference between the levels of mRNAs and proteins suggests the presence of additional post-transcriptional regulatory mechanisms, and underscores the importance of validating transcriptomic data with protein-level analysis for accurate cell population characterization. Next, we investigated protein expression in glial cells and their spatial relationship with neurons. SGCs in humans express the markers Glial fibrillary acidic protein (GFAP) and Glutamine synthetase(*11*), although GFAP is expressed at low level in normal condition and is upregulated under pathological conditions in rodents (*68*). We observed that GFAP protein was strongly enriched in SGCs, which formed a characteristic circular pattern around neuronal soma, even though *GFAP* transcripts were barely detected (**Fig. 4C, D),** consistent with our previous studies(*17*). We noted that some of the GFAP positive SGCs also extended to the proximal axon segment, thereby covering both the soma and axon initial segment, consistent with the notion that SGCs can ensheathe the initial region of the axon, known as Cajal’s initial glomerulus (*69*)(**Fig 4D**). In close contact with the SGCs, we identified a second glial population characterized by the expression of SCN7A. These SCN7A positive cells were positioned around the soma, forming a second larger ring, and also extended along the axon (**Fig 4D**). In the axon rich areas of the DRG, we observed both nmSCs (SCN7A positive) and mySCs, marked by *MBP* expression (**Fig. 4E, F)**. These findings demonstrate a layered organization of two distinct non-myelinating glial cell types around sensory neuron soma, with SGCs forming the innermost ring and another glia cell establishing close contact with both SGCs and axons. The precise nature of the outermost *SNC7A* positive glial cells surrounding neuronal soma and the spatial organization of the transcriptionally distinct nmSC_1 and nmSC_2 will require further studies.

We then examined the immune cell population, which are known to contribute to pain(*6*), repair(*70, 71*), and also monitor the vasculature in the DRG(*72*), at least in rodents. We identified three subpopulations of macrophages, all of which had highly enriched expression of Spleen Associated Tyrosine Kinase (*SYK*) (**Fig 3B**), a critical regulator of macrophage-mediated inflammatory responses(*56*). Mac_*MRC1* also had a highly unique expression of *F13A1, MRC1, LYVE1, CD163*, and *SIGLEC1*, genes characteristic of brain perivascular macrophages(*73*). CD163+MRC1+ macrophages are transcriptionally, anatomically, and ontogenically conserved between human and mouse DRGs(*72*). They are self-maintained locally, specifically involved in vasculature monitoring, and displayed distinct responses during peripheral inflammation(*72*). Mac_TOP2A was named because of the unique expression of multiple proliferation markers such as *TOP2A* and *MKI67* and there is evidence in rodents that macrophages in DRG have the capacity to self-renew (*71*). The ontogeny and functions of Mac_*OLR1* remained to be studied. We also identified two fibroblast/mesenchymal cell populations characterized by high expression of conventional fibroblast marker including *EGFR, APOD, DCN, PDGFRA, and COL1A1. The Fibro_Epi population* were uniquely enriched for *FBLN1 and EYA1, marker genes for epineurium fibroblast whereas Fibro_Endo+Peri were enriched for known endoneurium and perineurium fibroblast markers*. Although recent studies have shown that the DRG fibroblasts regulate neuropathic and mechanical nociception in mice(*74, 75*), the underlying molecular mechanisms remain largely unknown.

### Mapping and characterization of human DRG open chromatin landscape

To define the gene-regulatory programs underlying the identity and function of each DRG cell type, we identified the open chromatin regions in each of the 14 DRG cell types. Since peak calling based on the annotated cell clusters enables the identification of more cell-type-specific peaks(*53, 76*), we therefore aggregated the chromatin accessibility profiles from the nuclei comprising each cell cluster and identified the cell-type-specific open chromatin regions with MACS2 (*50*) using a previously reported approach(*77*). This resulted in a uniform peak set of 501 bp which were used for all downstream analyses. We detected a total of 278,208 open chromatin regions (referred to as OCRs) across all 14 cell types (**Fig. S4A**), with SGCs having the most OCRs. We detected very few OCRs for the Neuron_2 clusters likely due to their small number and highly heterogenous cell content. Of all OCRs, 15.75% were located at annotated promoter regions, 48.75% in introns, 25.75% in the distal intergenic regions (**Fig. S4B**). The OCRs for canonical marker genes are consistent with the marker gene expression of major cell-type annotations (**Fig. 3C**). Each cell type was characterized by many specifically expressed genes and accessible OCRs (**Fig. S4C–D**).

### Transcriptional regulatory network governing human DRG cell type identities

Genomic regulatory elements such as enhancers, most of which lie in non-coding regions of the genome, control gene expression in a cell-type and tissue-specific manner. To investigate transcriptional regulatory mechanisms underlying cell-type–specific gene expression in hDRG, we applied the computational framework FigR(*78*) to link distal cis-regulatory elements with target genes and infer gene-regulatory networks (GRNs) governing cell identity and function (**Fig. 5A**). Because chromatin accessibility and gene expression were simultaneously profiled in the same cells, we performed an integrative analysis using both snATAC-seq and snRNA-seq data. FigR was first used to identify significant distal peak-to-gene expression interactions by correlating accessibility of peaks within a fixed 100 kb window around each transcription start site (TSS) with expression of that gene, followed by permutation-based testing. We identified a total of 40,607 unique chromatin accessibility peaks significantly associated with gene expression (permutation p ≤ 0.05), spanning 12,401 genes within 100 kb of each other. Applying the default parameter of n ≥ 7 significant peak-gene associations per gene, we identified genes with a high density of peak-gene interactions, referred to as “domains of regulatory chromatin” (DORCs)”(*78*). In total, we found 1,532 DORC genes associated with 14,548 total peaks (**Fig. 5B, Table S6**).

**Figure 5:**
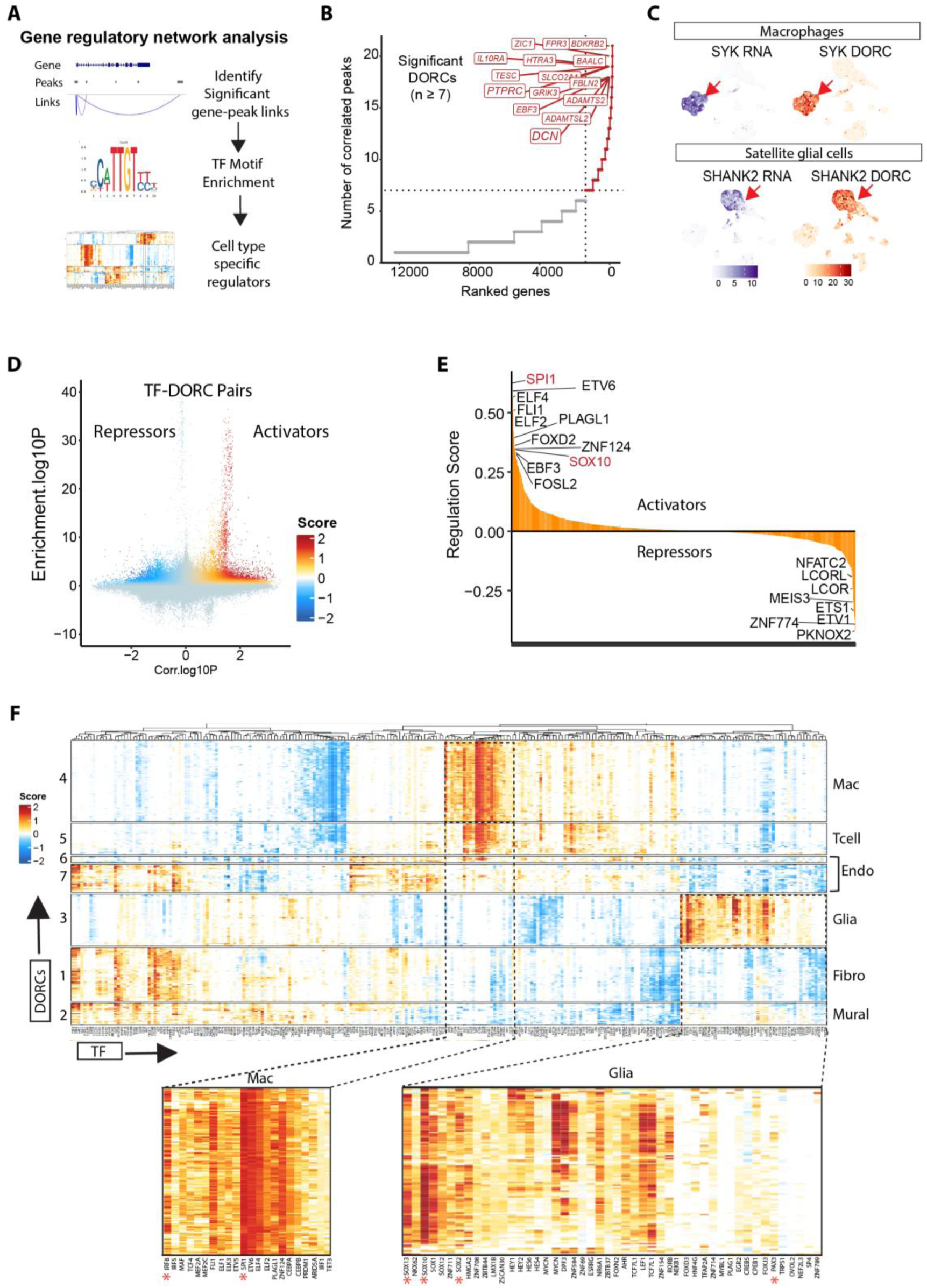
Integrated transcriptional regulatory network analysis identifies key regulatory modules for each cell type. (**A**) Schematic of transcriptional regulatory network analysis. (**B**) Significant DORCs identified by FigR. (**C**) UMAP of DORC accessibility scores (right) and paired RNA expression (left) for macrophage marker gene SYK and SGC marker SHANK2. (**D**) Scatterplot showing all DORC-to-TF associations, colored by the signed regulation score. (**E**) Mean regulation score (signed, −log10 scale) across all DORCs (n = 1,532) per TF, highlighting select TF activators (left skewed) versus TF repressors (right skewed). (**F**) Consensus k-mean clustering heatmap of DORC regulation scores for all significant DORC-to-TF associations. Rows indicate DORCs. Columns indicate TFs. The regulatory modules for macrophage and glia cells were enlarged at the bottom of the figure.

The list of DORC genes encompassed many known cell-type specific marker genes, including *FLT1* and *ERG* for endothelial cells; *DCN, APOD, EGFR, PDGFRA, FBLN1* for fibroblasts; *SYK, PTPRC, BCL11B, CD2, CSF1R, MRC1, F13A1, LYVE1,* and *PLAC8* for immune cells; NCMAP for mySC, *SCN7A, SEMA3E, and SORCS3* for nmSC; *MYOCD, RGS5* for mural cells; as well as *CTTND2* and *SHANK2* for SGC (**Fig. 5C, S5**). The strong correspondence between DORC accessibility scores and RNA expression levels in the same cell populations suggest that FigR successfully captured the genes and regulatory sequences defining each cell type.

Next, we applied FigR to identify putative transcription factor (TFs) regulators (both activators and repressors) of DORCs. Specifically, we examined the enrichment of different TF binding motifs within DORCs and the correlation of TF expression levels with the overall accessibility level of each DORC gene (DORC score). FigR integrates these two dimensions to compute a signed probability score called the “regulation score” (on a −log10 scale), which represents the regulatory activities of motif-enriched and RNA-correlated TFs. Positive values indicate activators and negative values indicate repressors. This analysis generated 526,084 TF-DORC associations (all tested TFs across all tested DORCs), with each point representing one TF-DORC pair (**Fig. 5D**). We identified 871 significant regulators across all cells for the given DORCs by filtering TF-DORC associations using a threshold of absolute (regulation score) ≥ 1(*78*) (**Table S7)**. Ranking TFs by their mean regulation score across DORCs, among the top ranked activators were well-established cell identity regulators, such as *SPI1* (PU.1) for macrophage (*79*) and *SOX10,* which is required for specification of peripheral nervous system (PNS) glial cells (*80*) (**Fig. 5E**).

To uncover cooperative TFs networks, we performed consensus k-means clustering on the TF-DORC regulation scores. This analysis grouped TFs (column) that may coordinately regulate modules of DORCs (rows) (**Fig. 5F).** Notably, we identified transcription regulatory networks underlying cell-type-specific gene signature modules, including glia (module 3 :*CTNND2, SHANK2, SORCS3*), macrophage (module 4: *SYK, CLEC7A, C3, IRF8, PLAC8*), T cells (module 5: *PTPRC, CD69, STAT4*), endothelial cells (module 6 and 7: *FLT1*), and fibroblasts (module 1: *DCN, APOD, FBLN1, EGFR*), and mural cells (module 2: *EDNRA, NOTCH3*) **(Table S8)**. The predicted TFs for each cell-type-specific gene module aligned well with established biology. For example, *SPI1* (PU.1), *JUNB, IRF8, RUNX1, MEF2A, and MEF2C* are key regulator of macrophage identity(*81*), while *SOX10, SOX2*, *SOX5, SOX6,* and *PAX3* play critical roles in glia development and specification (*80*).

### SOX10 regulatory network in peripheral glial cells

Our gene regulatory network analysis identified *SOX10* as a top transcription activator regulating glial signature gene modules (**Fig. 5E, E)**, with predicted binding to 103 DORCs (**Table S9)**. These predicted targets included established glial markers and genes important for glial development and function in the PNS, such as MBP, *SORCS3, SHANK2, SOX1, SOX2, CTNND2, and PAX3* (*82–85*). To understand cell-type-specific functions of SOX10, we examined SOX10 expression and regulatory activity in individual cell types. While *SOX10* was expressed in neurons, mySC, nmSC, and SGC in the snRNA-seq data (**Fig. 6A**), TF foot printing analysis of SOX10 binding sites (**Fig. 6B**) revealed SGC, nmSC_1, and nmSC_2 as the top three cell populations, suggesting that SOX10 exhibit strong regulatory activities in glial cells (**Fig. 6C**). To examine SOX10 expression at the protein level, we performed immunofluorescence staining of SOX10 and co-labeled with GFAP, SCN7A, and MBP. SOX10 was localized in nuclei as expected for a TF, and these SOX10-positive nuclei localized both around neuronal soma and within axon-rich regions (**Fig. 6D, E)**. Around neuronal soma, SOX10 nuclei co-localized with GFAP in SGCs and with SCN7A positive cells (**Fig. 6D)**. In axon-rich areas, SOX10 colocalized with MBP in mySCs and with SCN7A in nmSCs (**Fig. 6E)**. Notably, SOX10 protein was absent from neurons, despite detectable transcripts in the snRNA-seq data (**Fig. 6A**), representing another example of a discrepancy between transcript and protein levels.

**Figure 6:**
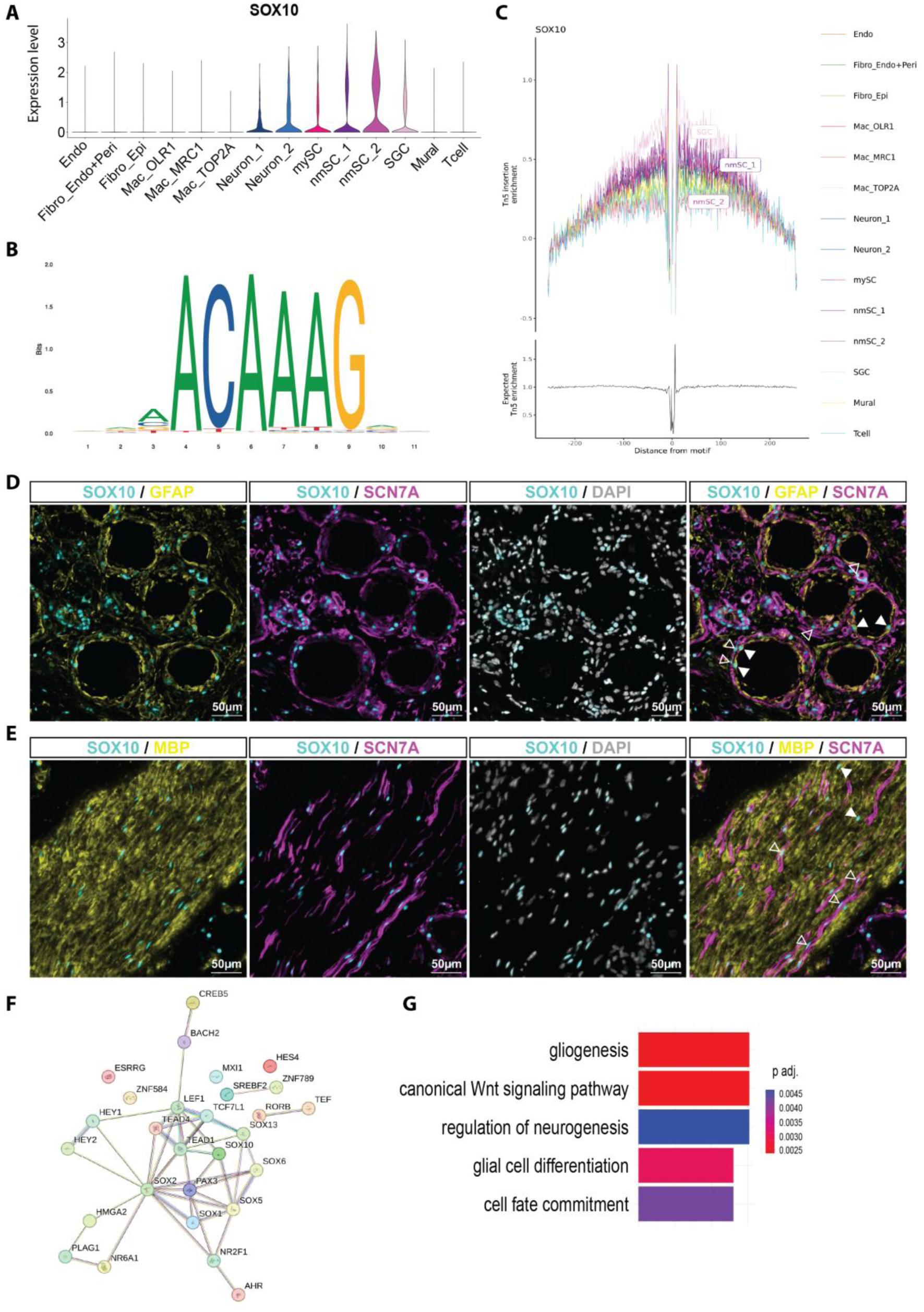
SOX10 regulatory network. (**A**) Violin-plots showing the expression of *SOX10* across the different cell populations. (**B**) Sequence logo for DNA sequence motif of SOX10. (**C**) TF footprinting analysis for SOX10 motifs sites. (**d, e**) Representative images of immunofluorescence staining of neuron soma dense area (D) and axon bundles dense area (e) of hDRG, labeled for SOX10 (cyan), SCN7A (magenta), DAPI (gray), and GFAP (D) or MBP (E) in yellow. (**D**) Solid arrows indicate SOX10-positive nuclei in GFAP-enriched cells. Hollow arrows indicate SOX10-positive nuclei in SCN7A-enriched cells. (**E**) Solid arrows indicate SOX10-positive nuclei in MBP-enriched cells. Hollow arrows indicate SOX10-positive nuclei in SCN7A-enriched cells. (**F**) Protein-protein interaction network for the candidate TFs that regulate glia DORCs. (**G**) Gene ontology terms enriched in the candidate TFs that regulate glia DORCs. Scale bar: 50 µm.

Our GRN analysis revealed that binding sites for many TFs were concordantly enriched in the glial DORC modules (**Fig. 5F**), suggesting cooperative regulation of glial identities and functions. These included multiple high mobility group (HMG) domain transcription factors—*SOX10, SOX2*, *SOX5, SOX6,* and *PAX3*, which can interact with each other through protein-protein interaction as revealed by the String database (**Fig. 6F**). Sox10 and Pax3 physically interact with each other to activate c-RET in neural crest cells in mice, which is critical for enteric ganglia development(*86*). Similarly, Sox10, Sox5 and Sox6 act synergistically to regulate oligodendrocyte maturation in the mouse central nervous system (CNS) (*87, 88*). Consistent with their role in glia fate specification during development in the CNS, these TFs were significantly enriched for pathways including “Cell fate commitment” and “Glial cell differentiation” (**Fig. 6G**). Our analysis suggests that these TFs may cooperate with each other and play critical roles in defining glial cell identity and function in the hDRG.

Similarly, SPI1 was identified as the top-ranked transcription activator in macrophages (**Fig. 5E, Table S7**). SPI1 expression was restricted to macrophage populations (**Fig. S6a**), and TF foot printing analysis identified Mac_OLR1, Mac_MRC1, and Mac_TOP2A as the top cell populations most enriched for SPI1 binding sites (**Fig. S6B**). We identified 241 putative SPI1 target DORCs, including common macrophage marker genes such as *PTPRC*, *CLEC7A*, *CSF1R*, *F13A1*, *IRF8*, *PLAC8*, *SYK*, and *C3* (**Fig. S6A**, **Table S10**), consistent with SPI1’s established role in macrophage cell fate regulation (*89, 90*). IRF8 has been shown to be critical for transforming microglia (the resident macrophage of the brain and spinal cord) into a reactive phenotype, which is associated with neuropathic pain after peripheral nerve injury in mice(*91*). Furthermore, SPI1 and IRF8 together regulate cell type-specific expression of NLRP3 inflammasome in human macrophages(*92*), consistent with our GRN analysis that binding sites for these two TFs are concordantly enriched in the macrophage DORC modules (**Fig. 5F**).

### Defining cell types and putative regulatory mechanisms critical for pain conditions

Genome-wide association studies (GWAS) have successfully identified many disease-associated loci. However, most GWAS studies are performed on bulk tissue samples, representing complex mixtures of cell types, whereas disease-causal variants often act in specific tissues and cell types. Therefore, the causal variants, their target genes, and the relevant cell types largely remain unsolved. Single-cell multi-omic datasets now offer a powerful opportunity to predict disease-relevant cell types and pinpoint causal variants by jointly examining gene expression and chromatin accessibility. We first curated a comprehensive set of pain-associated SNPs and genes. In total, we collected 6,212 SNPs across seven published GWAS studies of migraine and multiple pain conditions, along with their associated genes (**Table S11**). The following studies were included: 1) Gormley et al.(*93*) identified 38 migraine susceptibility loci (referred to as Migraine_PG) through a meta-analysis of 375,000 individuals from the International Headache Genetics Consortium. 2) Hautakangas et al. (*94*) performed a GWAS of 102,084 migraine cases and identified 123 risk loci and subtype-specific risk alleles (Migraine_HH). 3) Toikumo et al. conducted a cross-ancestry meta-analysis of pain intensity in 598,339 participants in the Million Veteran Program, identifying 126 independent genetic loci, including 69 novel ones (Painintensity_ST). 4) Khoury et al. (*95*) reported 896 SNPs spanning 23 loci of chronic multi-site pain (Chronic_Multi_SK) in the UK Biobank. Although they did not find replicated SNPs associated with chronic single-site pain we included their reported single-site SNPs in our analysis (Chronic_Single_SK). 5) Johnston et al.(*26*) identified 39 SNPs associated with multi-site chronic pain in the UK Biobank (Chronic_Multi_KJAJ). 6) Zorina-Lichtenwalter et al. (*34*) identified 315 common genetic factors shared across 24 distinct pain conditions in over 400,000 individuals in the UK Biobank participants (Chronic_Multi_KZL). 7) Wistrom et al. (*96*) compiled 242 genes linked to pain-associated behaviors through literature search spanning over three decades (Pain_EW). 8) Lastly, we included 28 genes (Neuropathy) associated with hereditary sensory and autonomic neuropathies (HSAN) provided at the HEREDITARY SENSORY & AUTONOMIC NEUROPATHIES (HSAN) website (https://neuromuscular.wustl.edu/time/hsn.htm).

We then used snRNA-seq data to identify disease-relevant cell types in the hDRG. Using cell cluster-specific marker genes, we performed hypergeometric tests to identify cell populations that were significantly enriched for pain-associated gene sets (**Fig. 7A**). Neuron_1 marker genes were significantly enriched across nearly all pain conditions (except migraine), consistent with the finding that Neuron_1 includes nociceptor populations. Neurons in trigeminal ganglia, which have a different developmental origin compared to DRG neurons, might be more relevant to migraine. Interestingly, Mac_OLR1 marker genes were significantly enriched for all pain conditions, further supporting the critical role of macrophages in pain processing (*6, 70*). Fibroblast, mural cells, endothelial cell and glial cell marker genes were also enriched in subsets of pain conditions, highlighting that non-neuronal cells play critical roles in pain pathogenesis.

**Figure 7:**
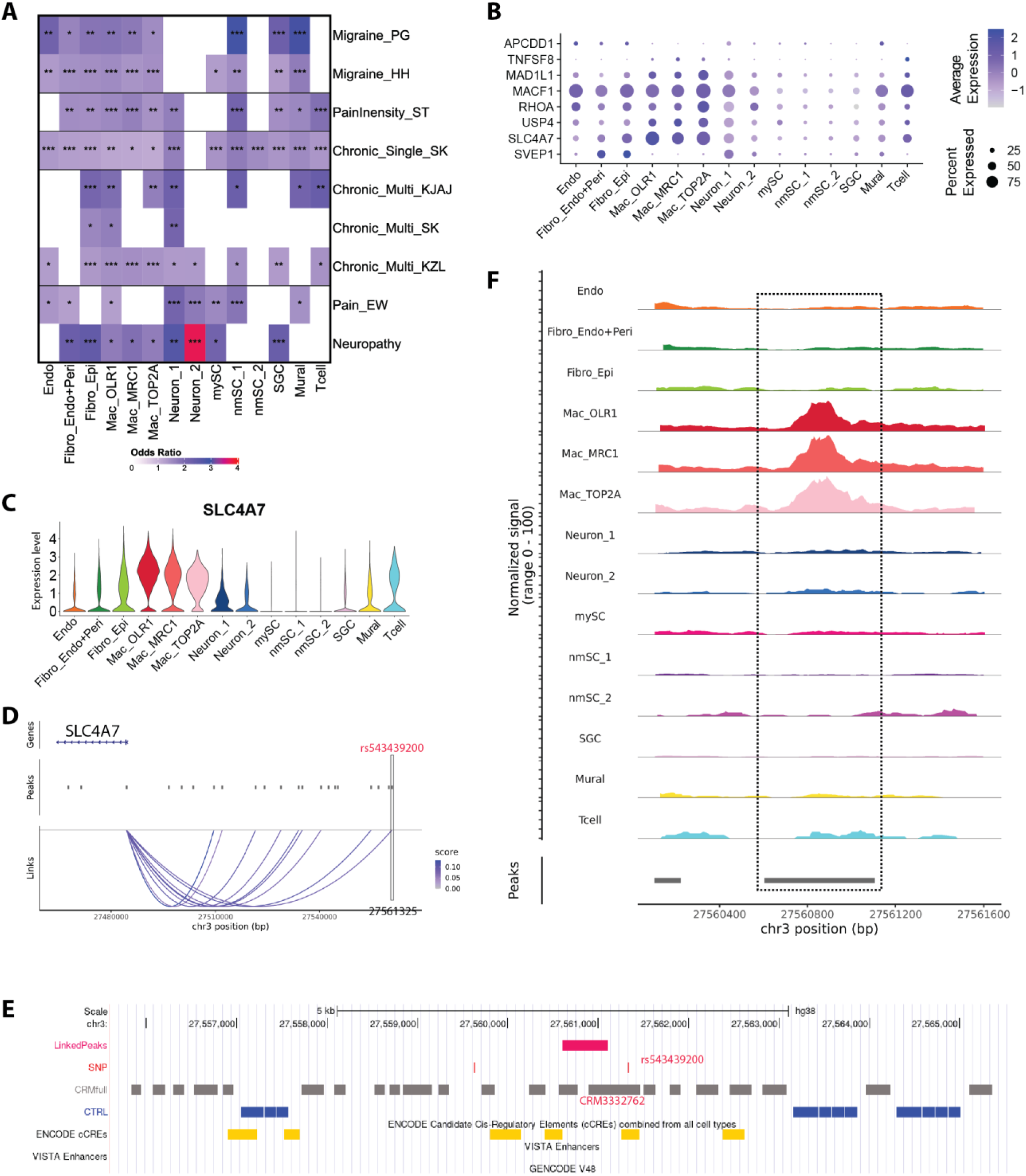
Defining cell types and putative regulatory mechanisms critical for pain conditions. (**A**) Heatmap showing cell types whose marker genes significantly enriched for pain-associated genes. (**B**) Dot plot showing relative expression of all pain-associated genes significantly linked to snATAC-seq peaks. (**C**) Violin-plot showing the expression levels of *SLC4A7* across all cell populations. (**D**) peak–gene links for *SLC4A7* and the location of pain-associated SNP rs543439200. (**E**) Genomic view of the snATAC-seq peak significantly associated with *SLC4A7*, the location of pain-associated SNP rs543439200, VRMOD predicted CRM3332762, and ENCODE cCREs. (**F**) snATAC-seq peak signals.

Next, we used snATAC-seq data to identify putative causal variants through a disease SNP prioritization framework (*97*). This framework integrates cis-regulatory modules (CRMs) predicted by the Vertebrate Regulatory MOdule Detector (VRMOD) with ten orthogonal human datasets of experimentally defined CRMs as well as genomic and epigenomic datasets (details in Method). Gene transcription is regulated by TFs binding to cognate transcription factor binding sites (TFBSs), which often cluster into CRMs, to ensure precise spatiotemporal gene expression(*98, 99*). Importantly, GWAS-reported SNPs are usually tag-SNPs—representative variants that capture the signal of nearby correlated SNPs through linkage disequilibrium.

Because tag-SNPs may not be causal, it is essential to prioritize functional non-coding variants located within regulatory modules. VRMOD accurately predicts functional CRMs directly from genomic sequences alone. CRMs supported by both VRMOD predictions and orthogonal experimental datasets represent the strongest candidates for harboring transcriptional regulatory activities. For integration, we included ten human experimental datasets including experimentally defined regulatory modules as well as genomic and epigenomic datasets representing putative regulatory sequences from six sources including enhancers from VISTA Enhancer Browser (*100*), predicted enhancers from the Enhancer Atlas(*101*), Chromatin accessibility (ATAC-seq and DNAse-seq) peaks, histone modification ChIP-seq peaks (H3K27ac, H3K4me1, and H3K4me3), ENCODE candidate cis-regulatory elements (cCREs)(*102*), and scATAC-seq peaks from 45 adult and fetal human tissues(*103*). By integrating GWAS pain-associated SNPs with our single cell ATAC-seq data through this framework, we prioritized genomic regions most likely to harbor true functional non-coding variants that are physically linked to GWAS tag-SNPs within the same CRM.

We first identified VRMOD-predicted CRMs that contained at least one pain-associated SNP (hereafter referred to as pain-associated CRMs). We then linked these CRMs to potential target genes by overlapping them with peaks significantly associated with genes located within 100kb of a transcription start site (TSS), based on the genome wide peak-gene linkage analysis described above. In total, we found eleven peaks that overlapped (≥ 1bp) with ten pain-associated CRMs. These peaks were linked to eight unique genes: *SVEP1, SLC4A7, USP4, RHOA, MACF1, MAD1L1, TNFSF8, APCDD1* (**Fig. 7B, Table S12**). Although these genes were expressed across all cell types, they were generally de-enriched in glial cells, with some showing cell-type-specific enrichment in other cell types. Three of these genes (*SLC4A7*, *RHOA* and *USP4*) have known association with pain. *SLC4A7* was identified as a gene associated with a single-site chronic pain SNP(*95*). *SLC4A7* is strongly enriched in macrophages (**Fig. 7B, C**) and encodes the electroneutral sodium/HCO3 cotransporter NBCn1, which plays important physiological and pathophysiological roles in diverse cell types(*104*). Located on human chromosome 3, *SLC4A7* had 12 significantly associated peaks within 100 kb of its TSS (**Fig. 7D**), suggesting complex transcriptional regulation. We identified a VRMOD predicted module, CRM3332762, that encompassed the pain-associated SNP rs543539200 and overlapped with both an ATAC-seq peak (chr3:27560605-27561105) and an ENCODE cCRE (**Fig. 7E**). These lines of evidence support the role of CRM3332762 as a regulatory element modulating *SLC4A7* expression and potentially rs543539200 as a causal variant contributing to pain conditions. Interestingly, the ATAC-seq peak chr3:27560605-27561105 overlapping CRM3332762 was unique to macrophages, further suggesting that SNP rs543539200 may regulate *SLC4A7* expression in a macrophage-specific manner.

Both Ras homolog gene family member A (*RHOA*) and Ubiquitin-specific protease 4 (*USP4*) were associated with pain-linked SNP rs4955440, a multi-site chronic pain SNP located between the two genes (*95*). Both genes were broadly expressed across nearly all cell types, with varying expression levels, except nmSCs (**Fig. 8A, B**). We identified two significant peaks associated with *RHOA* and three with *USP4* within 100kb of their respective TSS (**Fig. 8C**). The pain-associated SNP rs4955440 was located within VRMOD-predicted CRM3377162, which overlapped with an snATAC-seq peak (chr3:49350847-49351347), and an ENCODE cCRE (**Fig. 8D**), supporting its putative regulatory function. Rho-associated coiled-coil-containing protein kinase (ROCK) is the primary downstream effector of RhoA(*105*). Dysregulated activation of the RhoA/ROCK signaling pathway has been reported in the DRG and spinal cord in rodent pain models, and pharmacological inhibition of this pathway alleviates hyperalgesia and allodynia(*106*). *USP4,* in contrast, exhibits context-dependent roles in injury and inflammation. Following spinal cord injury, *USP4* expression is downregulated, promoting microglial activation and subsequent neuronal inflammation(*107*). Conversely, in a liver injury model, *USP4* levels were elevated, where it functions to reduce hepatocyte apoptosis and liver inflammation(*108*). These findings underscore the cell-type- and context-specific regulation of USP4. Interestingly, the snATAC-seq peak chr3:49350847-49351347, overlapping CRM3377162, was unique to macrophages (**Fig. 8E**), suggesting a potential macrophage-specific regulatory role of rs4955440 in pain susceptibility.

**Figure 8:**
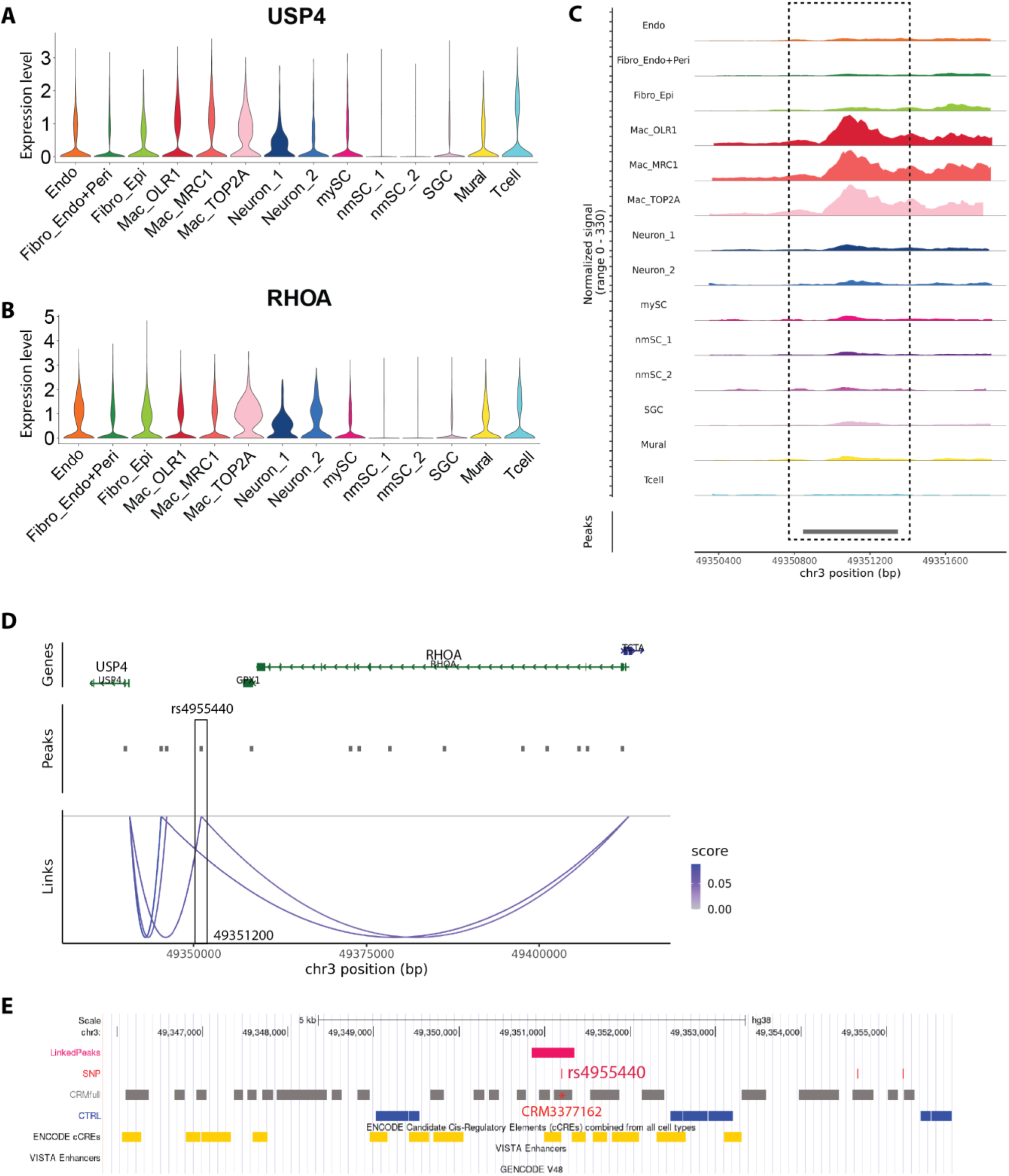
Macrophage-specific association of pain-associated SNP rs4955440 linked to *RHOA* and *USP4*. (**A, B**) Violin-plots showing the expression of *RHOA* (A) and *USP4* (B) across different cell populations. (**C**) Peak–gene links for *RHOA* and *USP4* and the location of pain-associated SNP rs4955440. (**D**) Genomic view of the snATAC-seq peak significantly associated with *RHOA* and *USP4*, the location of pain-associated SNP rs4955440, VRMOD predicted CRM3377162, and ENCODE cCREs. (**F**) snATAC-seq peak signals.

We also identified five new putative pain-associated genes, *SVEP1, MACF1, MAD1L1, TNFSF8, APCDD1*. *SVEP1* is an extracellular matrix protein that causally promotes vascular disease(*109*) and associates with multiple other diseases including hypertension, type 2 diabetes, glaucoma(*110, 111*), and neuropathic pain(*112*). *SVEP1* is uniquely expressed in fibroblasts and Neuron_2 (**Fig. S7A).** Interestingly, many pain-associated SNPs were located within the first intron of *SVEP1* (**Fig. S7C**). The first introns of higher eukaryotes are typically longer and evolutionary conserved than other introns and positively associated with regulatory epigenetic signals(*113*). However, within the first intron, only two SNPs -- rs77748855 and rs80253943(*95*), were located within a VRMOD-predicted CRM (CRM5626436) that also overlapped with snATAC-seq peak (chr9:110575075-110575575), suggesting potential regulatory activity in DRG cell types. This snATAC-seq peak exhibited strong chromatin accessibility in multiple cell types including endothelial cells, both fibroblast populations, both neuronal populations, and nmSC cells. Notably, robust *SVEP1* transcript expression was detected only in the two fibroblast populations and in Neuron_2, but not in the other cell types (**Fig. S7D**), indicating cell-type-specific regulatory functions of CRM5626436 and the two SNPs. *MAD1L1* encodes the spindle assembly checkpoint (SAC) protein MAD1, which is an essential component of the mitotic checkpoint. *MAD1L1* mutations cause a new variant of mosaic variegated aneuploidy with systemic inflammation and unprecedented tumor susceptibility(*114*). Whether *SVEP1* or *MAD1L1* play any role in pain sensation remains to be tested.

## DISCUSSION

The mechanisms underlying chronic pain in various conditions are not fully understood, and existing treatments often prove ineffective for most patients(*30–34*). There is thus an urgent need for advancements in our understanding of the mechanisms regulating pain transmission, which could generate new pivotal insights for the treatment of chronic pain. DRGs harbor the soma of sensory neurons, which integrate signals from the body’s internal and external environment, representing a first step in pain signal transduction. Our study provides a deeper understanding of the transcriptional and regulatory landscapes within all cell types in hDRG, emphasizing the critical roles played by non-neuronal cell types, such as endothelial cells, fibroblasts, macrophages, and glial cells in the pathogenesis of pain. Furthermore, our study underscores the importance of sequencing depth in enhancing data resolution and accuracy, which in turn facilitates refined cell-type clustering and more precise scientific interpretations. Our study also highlights the need for cell-type validation at the protein level, because of the weak correlation between gene and protein expression.

Due to considerable differences between animal models and humans, there is an urgent need to elucidate the molecular profile of human sensory neurons and non-neuronal cells to develop meaningful strategies to treat pain. With the ability to acquire and study hDRG through organ procurement organizations, the last decade has seen significant advances in characterizing the cellular and molecular features of cells in hDRG (*17, 20, 21, 23, 25, 112, 115–120*). Most efforts have focused on transcriptional profiling to identify neuron and non-neuronal cell types, determine interspecies similarities and differences, and probe mechanisms underlying acute and chronic pain(*17, 20, 21, 115, 116*). Here, we extend beyond transcriptomics to examine the chromatin landscape within the same cells. By integrating the transcriptomic and epigenetic data from single-cell multi-omics analysis of hDRG with pain-associated genes and SNPs identified in multiple GWAS, we infer susceptible cell types and candidate causal variants underlying human pain disorders. Our analysis uncovered three known pain-causal genes, five new putative pain-associated genes, and the cell-type-specific regulatory mechanisms that may underline their contributions to pain.

The large size of human neurons and their low abundance within DRG tissue represent a major obstacle to defining their molecular profiles. Our results further emphasize that even with single-nuclei sequencing, which circumvents challenges posed by the large neuronal soma in microfluidic-based single-cell RNA-seq technologies, neuronal recovery was limited, yielding only two broad populations. These findings highlight the need for profiling substantially larger cohorts of hDRG to capture sufficient neuronal diversity and enable accurate molecular characterization. Nonetheless, by leveraging a curated comprehensive set of pain-associated SNPs and genes, we found that cell types contributing to Neuron_1, but not Neuron_2, express marker genes significantly enriched across nearly all pain conditions (with the exception of migraine, as expected). This enrichment is consistent with the notion that Neuron_1 includes nociceptors.

Multiple non-neuronal cell types have emerged to contribute to pain development in animal models(*2–8*), underscoring the importance of characterizing the molecular profiles of non-neuronal cells in hDRG. Supporting this, our analysis of pain-associated SNPs derived from GWAS studies of various pain conditions revealed enrichment not only for neuronal marker genes, but also for marker genes of glial cells in subsets of pain conditions. We identified the three main glial types expected in DRG, namely SGCs, mySCs and nmSCs. Our gene regulatory network analysis identified SOX10 as a transcription activator regulating glial signature gene modules, and SOX10 was expressed in all three glial cell types at both transcript and protein levels. Interestingly, glial cells labeled for SCN7A, occupied two distinct spatial domains within the hDRG. SCN7A positive cells were found around the axon as expected for nmSC, but also, and surprisingly, around SGCs. Whether the SCN7A positive cells around the neuronal soma represent a subtype of SGCs or a specialized nmSC remains to be established. Future studies will also be needed to determine if the two nmSC populations we identified by unbiased clustering correspond to these two spatially distinct SCN7A-positive subtypes.

Our analysis of SNPs within cis-regulatory modules provided further support for non-neuronal cells in pain, especially immune cells and fibroblasts. The role of immune cells in pain has been well established in rodent models(*121*), and our study highlights a potential role for macrophages in human pain conditions. Notably, *SLC4A7*, which is enriched in macrophages with additional expression in fibroblasts, was identified as a gene associated with a single-site chronic pain SNP (*95*). *SLC4A7* encodes the electroneutral sodium/HCO3 cotransporter NBCn1, which plays important physiological and pathophysiological roles in diverse cell types. In animal models, SLC4A7 may be linked to sensory dysfunctions. For example, genetic disruption of *Slc4a7/*NBCn1 in mice leads to pronounced loss of hair cells and degeneration of sensory receptors in the eye and inner ear causing blindness and deafness(*122, 123*). In macrophages, SLC4A7 expression increases during differentiation from monocytes and is important for proper phagosome function in macrophages(*124*). Together, these findings suggest that regulation of SLC4A7 expression may be critical for sensory function and pain-associated symptoms.

Fibroblasts in the DRG remain poorly characterized. However, emerging evidence suggests that extracellular matrix (ECM) produced in part by fibroblasts contribute to neuronal signaling in both mice and humans, and that dysregulation of ECM genes is associated with neuropathic pain(*75, 112*). We uncovered multiple pain-associated SNPs within the first intron of SVEP1, an ECM protein previously implicated in vascular and chronic diseases such as hypertension, type 2 diabetes, and glaucoma. SVEP1 expression was detected in fibroblasts and the Neuron_1 population, further highlighting the role of ECM in pain processing. Additionally, our analysis also points to MAD1L1, a gene involved in the spindle assembly checkpoint, where mutations lead to mosaic variegated aneuploidy featuring systemic inflammation and heightened tumor susceptibility(*114*). Whether SVEP1 and MAD1L1 directly contribute to pain signaling in the DRG remains to be determined. Together, these findings underscore the potential importance of these newly identified genes in pain pathways and advocate for further investigation into their roles, which may offer new therapeutic targets for pain management.

In summary, this work provides new insights into the transcriptional regulation of diverse hDRG cell types and demonstrates that the neuronal microenvironment in the DRG is a critical component in pain pathogenesis. Our findings highlight candidate cell types and putative molecular mechanisms underlying pain sensation, offering a foundation for future studies aimed at therapeutic development.

## MATERIALS AND METHODS

### Human tissue

Human DRG were obtained from organ donors with full legal consent for use of tissue in research and in compliance with procedures approved by Mid-America Transplant. The Human Research Protection Office at Washington University in St. Louis provided an institutional review board waiver.

### Tissue preparation and immunohistochemistry

For immunohistochemistry studies, after isolation of mouse DRG, the tissue was postfixed using 4% paraformaldehyde for 24 hours at 4 degrees. Tissue was then washed in PBS and cryoprotected using 30% sucrose solution at 4 °C overnight. Next, the human and mouse tissues were embedded in optimal cutting temperature, frozen, and mounted for cryosectioning. Human and mouse frozen sections were cut at 7 µm and 10 µm respectively mounted on Superfrost glass slides and stored at - 80°C. The day of the experiment, slides equilibrate at -20°C for 1h, then a 4°C for 10 minutes. Human sections were fixed with 4% paraformaldehyde for 30 minutes at 4 degrees. Sections were washed 3 times for 15 min each in 0.02 M PBS at room temperature. Block slide with 3% BSA, 0.3% Triton-X in DPBS for 45 min at room temperature. Sections were washed 3 times for 15 min each in 0.02 M PBS at room temperature. Human and mouse sections were incubated overnight in 0.02 M PBS containing 0.3% Triton X100, 0.5% BSA and primary antibodies at room temperature in a humid chamber. The following day, sections were washed 3 times for 15 min each in 0.02 M PBS and incubated for 2 h at room temperature with secondary antibodies, diluted in 0.02 M PBS containing 0.3% Triton X100 and 0.5% BSA. Then, sections were washed 3 times for 15 min in 0.02 M PBS and mounted using Vectashield medium containing DAPI (VectorLabs, H-2000-10). Images were acquired at 310 or 320 using a Nikon TE2000E inverted microscope. Stained sections with only secondary antibody were used as controls. Antibodies were as follows: NEFH antibody (BioTechne, NB300-217), NEFL (CST, 10768), GFAP (Addgene, 1934450), PRPH (Abcam, ab269861), ROBO2 (Biotechne, AF3147), COL IV (Abcam, ab6586), FABP7 (Biotechne, AF3166), NeuN (SYSY,266 011), SCN7A (Biotechne, NBP1-87075), MBP (Millipore Sigma, AB9348), SOX10 (BioTechne, AF2864).

### Neuron proportion quantification

To determine the percentage of neurons in adult human DRGs, we quantified the proportion of neurons in hDRG sections from three donors using pan-neuronal maker TUJ1 to label neurons, GFAP to label glial cells, and DAPI to label all nuclei. Due to the uneven distribution of neurons in the DRG, neuron counting was performed at two distinct magnifications. Low magnification (4X) was utilized to quantify the number of neurons in six to seven complete DRG sections per donor. Higher magnification (20X) facilitated the quantification of neurons in 12–15 neuron-dense areas per donor.

### snMulti-omic protocol comparison

We compared four nuclear dissociation methods previously described: a nuclear extraction protocol with density gradient centrifugation alone (Opti), Opti followed by FACS (Opti/FACS), a crude extraction protocol followed by FACS (Non-gradient/FACS), and the nuclear isolation kit developed by 10X Genomics (10X kit). Opti or Opti/FACS were preformed according to the protocol described previously with minor modifications(*125*). Human DRG sections were minced by scissors in homogenization buffer (0.25 M sucrose, 25 mM KCl, 5 mM MgCl_2_, 10 mM Tris-HCl, pH 8.0, 5 ug/mL actinomycin, 1% BSA, and 0.08 U/ul RNase inhibitor, 0.01% NP40) on ice. Samples were transferred to a dounce homogenizer for 15 strokes with the loose pestle in a total volume of 1 mL, followed by five additional 15 strokes with the tight pestle. The tissue homogenate was then passed through a 50 um filter and diluted 1:1 with working solution (50% iodixanol, 25 mM KCl, 5 mM MgCl_2_, and 10 mM Tris-HCl, pH 8.0). Nuclei were layered onto an iodixanol gradient after homogenization and ultracentrifuged as described previously. After ultracentrifugation, nuclei were collected between the 30 and 40% iodixanol layers and diluted with resuspension buffer (1xPBS with 1% BSA, and 0.08 U/ul RNase inhibitor). Nuclei were centrifuged at 500 g for 10 min at 4°C and resuspended in resuspension buffer. For Opti, nuclei were processed and loaded into microfluid chip according to the manual of 10X Genomics Multiome assay. For Opti/FACS, nuclei were stained with 7-AAD and 7-AAD+ events were sorted using 100 um nozzle on a BD FACSARIA II into a 1.5 mL microcentrifuge tube containing 100 ul of resuspension buffer. FACS-sorted nuclei were processed and loaded into microfluid chip according to the manual of 10X Genomics Multiome assay. For Non-gradient/FACS, human DRG sections were minced and homogenized as described in Opti. The tissue homogenate was then passed through a 50 um filter, and centrifuged at 500 g for 10 min at 4°C. Pellet was resuspended in resuspension buffer with 7-AAD. 7-AAD+ events were sorted using 100 um nozzle on a BD FACSARIA II into a 1.5 mL microcentrifuge tube containing 100 ul of resuspension buffer. FACS-sorted nuclei were processed and loaded into microfluid chip according to the manual of 10X Genomics Multiome assay. For 10X kit, nuclei were extracted following the manual of 10X Genomics Nuclear Isolation Kit and processed and loaded into microfluid chip according to the manual of 10X Genomics Multiome assay.

To assess the enrichment of neuronal nuclei, we performed snMultiome-seq on nuclei isolated by each protocol and sequenced the libraries with target sequencing depth at 50k reads per cell for the snRNA-seq library. We then performed clustering based on the snRNA-seq data using R package Seurat V4.3.0 and identified marker genes for each cluster. A cluster is identified as neuronal if nuclei in that cluster showed increased level of *RBFOX1* (avg_log2FC > 2, FDR < 0.01) compared to all other clusters. We then calculated the percentage of neuronal nuclei for each sequencing library. Opti yielded the most neuronal nuclei (2.39 ± 1.03%, n=4), while 10X, Opti/FACS, and non-gradient/FACS yielded 0.54 ± 0.07% (n=2), 0.06% and 0% neuronal nuclei, respectively. Opti-prep preserves neuronal nuclei, whereas neuronal nuclei are lost with the other methods. The Opti method (dx.doi.org/10.17504/protocols.io.5jyl8q4y8l2w/v1) was used for all additional samples. All samples were sequenced deeply targeting 150,000 reads per cell for snRNA-seq and 100,000 reads per cell for snATAC-seq.

### scRNA-seq data analysis

snRNA-seq data were analyzed using Seurat (v5.0.1). We assessed QC on a per sample basis before integration, uniformly removing all cells with nCount < 1000 and nFeature < 500, as well as cells with >10% of genes mapped to mitochondria. An additional cutoff for nCount was used to remove outlier cells which was determined by individual sample quality. We performed normalization using SCTransform on all samples, and feature selection and integration was performed per Seurat tutorial using default parameters. Dimensional reduction and cell clustering were performed using high dimensionality and resolution to identify low quality cells and doublets. After multiple rounds of iterative doublet/low quality cell removal and re-integration, a final object was generated for downstream analysis.

### snATAC-seq data analysis

snATAC-seq data were analyzed using Signac (v1.6.0). We calculated or obtained the following quality metrics from CellRanger ARC outputs: nucleosome signals, transcriptional start site (TSS) enrichment score, total number of fragments in peaks, fraction of fragments in peaks, and ratio reads in genomic blacklist regions. We removed cells with < 500 nFeatures and < 1000 nCounts, TSS enrichment score < 2, nucleosome signal > 2.

To generate uniform peak sets we first performed genome-wide chromatin accessibility peaks calling for each cell cluster using MACS v2 (MACS2), with the following flags explicitly set: --nomodel --call-summits --SPMR -q 0.01; generating a list of peak summit calls per cell clusters. Peak summits were then extended by 250 bases on either end to generate 501 bp peaks. The peaks for all cell clusters were then pooled and ranked by the FDR corrected p-values of the summit reported by MACS2. Overlapping peaks were iteratively removed keeping the peak with the lowest p-value until no overlapping peaks remained. Single cell counts for reads in peaks were generated by Signac as the total number of unique fragments cut sites overlapping a given peak window tallied for each unique cell barcode detected in the data, producing a matrix of single cell chromatin accessibility counts in peaks (rows) by cells (columns). This uniform peak set was then added to the final object and used for all downstream analyses. We performed normalization, feature selection, linear dimensional reduction, cell clustering following the tutorial of the Signac package. We create gene activity matrix, integrated with snRNA-seq data, and detected differentially accessible peaks between cell types as described in the Signac tutorial with default parameters.

### Multimodal Clustering

We performed a joint neighborhood analysis integrating the snRNA-seq data and snATAC-seq data using the weighted nearest neighbor (WNN) method in Seurat v5. We performed clustering using principal component analysis (PCA) on dimensions 1:30 for the RNA and dimensions 2:30 on the LSI components for the ATAC, followed by the functions “RunUMAP” and “FindClusters” with a resolution of 0.2. Cell types were then annotated using conserved and unique markers, as well as known markers with evidence in literature.

### Cluster marker genes and cluster open chromatin regions (referred to as OCRs)

Cluster marker genes and cluster open chromatin regions were calculated using the Seurat v5 “FindMarkers” function to identify transcripts or peaks present in at least 20% of cells with default parameters (an absolute log fold change ≥ 0.1, Bonferonni adjusted p-values *P*.adj < 0.05) to determine statistically significant genes and peaks.

### Computing peak-to-gene links

To calculate linkage scores between RNA transcripts and ATAC peaks, the “LinkPeaks” function from the Signac package was used. Peaks within 100kb of the TSS for all genes were evaluated and scored based on correlation between gene expression and chromatin accessibility. Significant peak-to-gene links were determined using default parameters for correlation coefficient (> 0.05) and p-value (< 0.05), as well as only considering links where the gene and peak are present in at least 10 cells.

### FigR gene regulatory network analysis

Peak-gene *cis*-regulatory correlation analysis was performed to determine high density domains of regulatory chromatin (DORCs) using FigR. The default 100 kb window around the TSS was used. The default parameter of n ≥ 7 significant peak-gene associations per gene was used to determine “domains of regulatory chromatin” (DORCs)”. A TF-DORC associations score threshold of absolute (regulation score) ≥ 1 was used to identify significant regulators as recommended.

### TF motif accessibility scoring

Single cell accessibility scores for TF motifs were computed using chromVAR(*126*) as previously described(*127*). For TF motif accessibility scores, the peak by TF motif overlap annotation matrix was generated using a list of human TF motif PWMs from the chromVARmotifs package in R (https://github.com/GreenleafLab/chromVARmotifs), and used along with the scATAC-seq reads in peak matrix to generate accessibility Z-scores across all scATAC-seq cells passing filter.

### Pain-associated cell types and putative GWAS causal variant identification

We integrated GWAS pain-associated SNPs together with our single cell ATAC-seq data through the VRMOD SNP and CRM prioritization framework. First, curated lists of SNPs and genes from GWAS for different pain conditions and diseases were collected from publications (**Table S11**). For SNPs where associated genes were not reported by the publication, we used ChipSeeker(*128*) to find the genes most closely associated with the reported SNPs. In total, we curated seven lists of SNPs from six publications with a total of 6,212 pain-associated SNPs.

Vertebrate Regulatory MOdule Detector (VRMOD)(*97*) relies solely on genomic sequences of a given organism for predicting cis-regulatory modules (CRMs), without using any empirical data or phylogenetic conservation information at any stage of CRM identification. As a result, any available empirical dataset can serve as orthogonal evidence, providing independent validation of predicted CRMs. For each CRM, we provide independent lines of evidence supporting its functionality through the integration of VRMOD prediction and ten genomic and epigenomic datasets representing putative regulatory sequences. The ten datasets are described below.

1. A total of 60 literature-curated human ExpCRMs driving gene expression in a diverse type of tissues, cell types, and developmental stages including neurons, glia and immune cells.
2. A total of 998 *in vivo* validated functional enhancer elements for human from the VISTA Enhancer Browser
3. A total of 201,032 putative human enhancers from Enhancer Atlas 2.0, annotated in 534 different tissues/cell types(*101*).
4. DHS peaks: DHS regions contain a variety of regulatory elements including enhancers, silencers, insulators, and locus control regions(*102*). Human DNase-seq data (2,401,388 peaks) were downloaded from the Cistrome database.
5. Bulk ATAC-seq peaks: The Cistrome database contained 1,217,002 human ATAC-seq peaks from 1,059 samples and 25 different tissues and cell types.
6. ChIP-seq data of the histone marks H3K4me1 and H3K27ac represent putative active enhancers. Human H3K4me1 (1,565,782) and H3K27ac (1,697,483) ChIP-seq data were downloaded from the Cistrome database.
7. ChIP-seq data of the H3K4me3, which represents putative promoters, were downloaded from the Cistrome database with 2,081,430 peaks for human.
8. scATAC-seq detects cell-type-specific open chromatin regions potentially representing cell-type-specific regulatory elements functioning in hundreds of cell types from different tissues and developmental stages. Human scATAC-seq data provided ∼1.2 million elements from 45 adult/fetal tissues and 222 cell types(*103*).
9. ENCODE cCREs are candidate cis-Regulatory Elements (cCREs) obtained through the integration of ENCODE epigenomics data. Currently there are two different versions of ENCODE cCREs. Our analysis was conducted before the release of ENCODE cCREs V3 (labeled as ENCODE cCREs herein). We used the ENCODE V2 version (referred to as ENCODE_V2 herein) to perform the analysis which included 1,063,878 cCREs from 1,518 tissues/cell types for the human genome(*102*).

To identify putative pain-causal SNPs, we converted pain-associated SNPs into bed format and identified VRMOD predicted CRMs that contain at least one pain-associated SNPs (referred to as **pain-associated CRMs**). Next, we identified genes potentially regulated by **pain-associated CRMs** using peaks that were significantly linked to genes within 100kb of gene TSSs determined via genome wide peak-gene link analysis.

Because these SNPs were located within cis-regulatory sequences with empirical data supporting their regulatory functions, they are likely SNPs that will affect target gene expression, therefore potentially causal to pain condition. We found ten unique peaks which overlapped with at least 1bp with ten pain-associated CRMs. Eight unique genes were significantly linked with these ten peaks.

## Acknowledgements

We thank all participants and their families for their commitment and dedication to advancing research of diagnosis and treatment for chronic pain, and the staff at Mid-America Transplant. We thank the Genome Technology Access Center at the McDonnell Genome Institute at Washington University School of Medicine for help with genomic analysis. The Center is partially supported by NCI Cancer Center Support Grant #P30 CA91842 to the Siteman Cancer Center from the National Center for Research Resources (NCRR), a component of the National Institutes of Health (NIH), and NIH Roadmap for Medical Research.

## Author Contributions

Conceptualization: KB, PM, LY, VC, GZ

Data generation: PM, LY

Methodology: KB, PM, LY, YC

Sample processing: AD, AC, BAC, JM, JDR, MP, PG, JY, RS, ZB, JL, SR

Investigation: KB, PM, LY, HN, GM Visualization: KB, PM, LY, HN

Funding acquisition: RWG, VC, GZ

Project administration: RWG, VC, GZ

Supervision: TW, RWG, VC, GZ

Writing – original draft: KB, PM, LY, HN, VC, GZ

Writing – review & editing: KB, PM, LY, RWG, VC, GZ, with feedback from all authors.

## Supplementary Materials

Fig S1 to S7 for multiple supplementary figures

Tables S1 to S12 for multiple supplementary tables

**Supplemental Table 1.** Donors’ metadata information.

**Supplemental Table 2.** Preliminary quantification of percentage of neuronal cells in human DRGs.

**Supplemental Table 3**. 10x Cellranger output statistics of shallow and deep sequencing data.

**Supplemental Table 4.** 10X CellRanger ARC output statistics of the 11 hDRG samples.

**Supplemental Table 5.** Top cluster marker genes.

**Supplemental Table 6**. DORCs identified by FigR analysis.

**Supplemental Table 7**. Transcription factors predicted by FigR.

**Supplemental Table 8**. Gene list for each regulatory module identified by consensus k-means clustering analysis.

**Supplemental Table 9.** Target genes of SOX10 predicted by FigR.

**Supplemental Table 10**. Target genes of SPI1 predicted by FigR.

**Supplemental Table 11**. Curated pain-associated SNPs and genes.

**Supplemental Table 12**. Putative pain-causal SNPs and linked genes.

---

**Figure S1.**
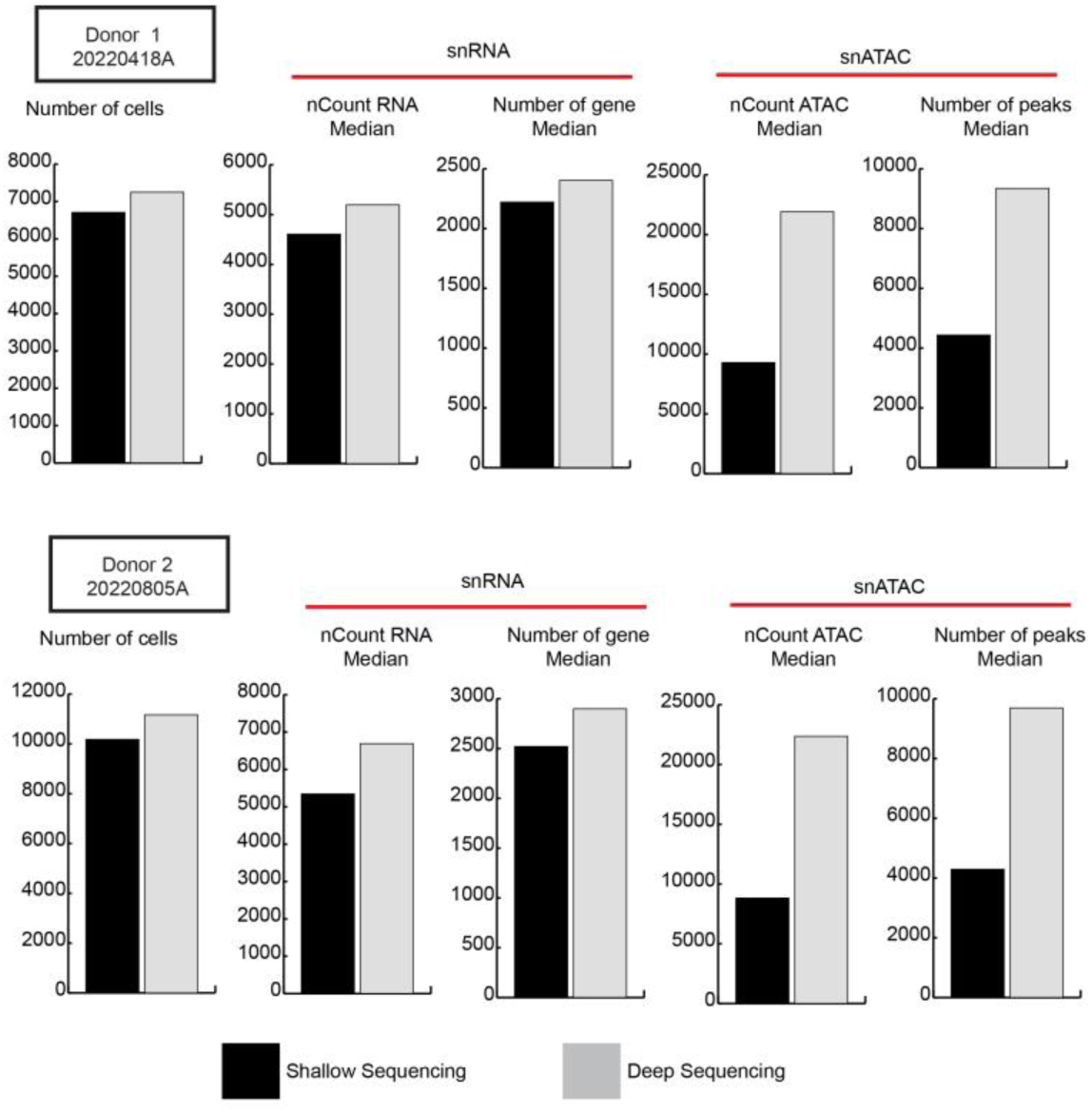
Sequencing statistics of shallow sequencing and deep sequencing data from Donor 1 and Donor 2.

**Figure S2.**
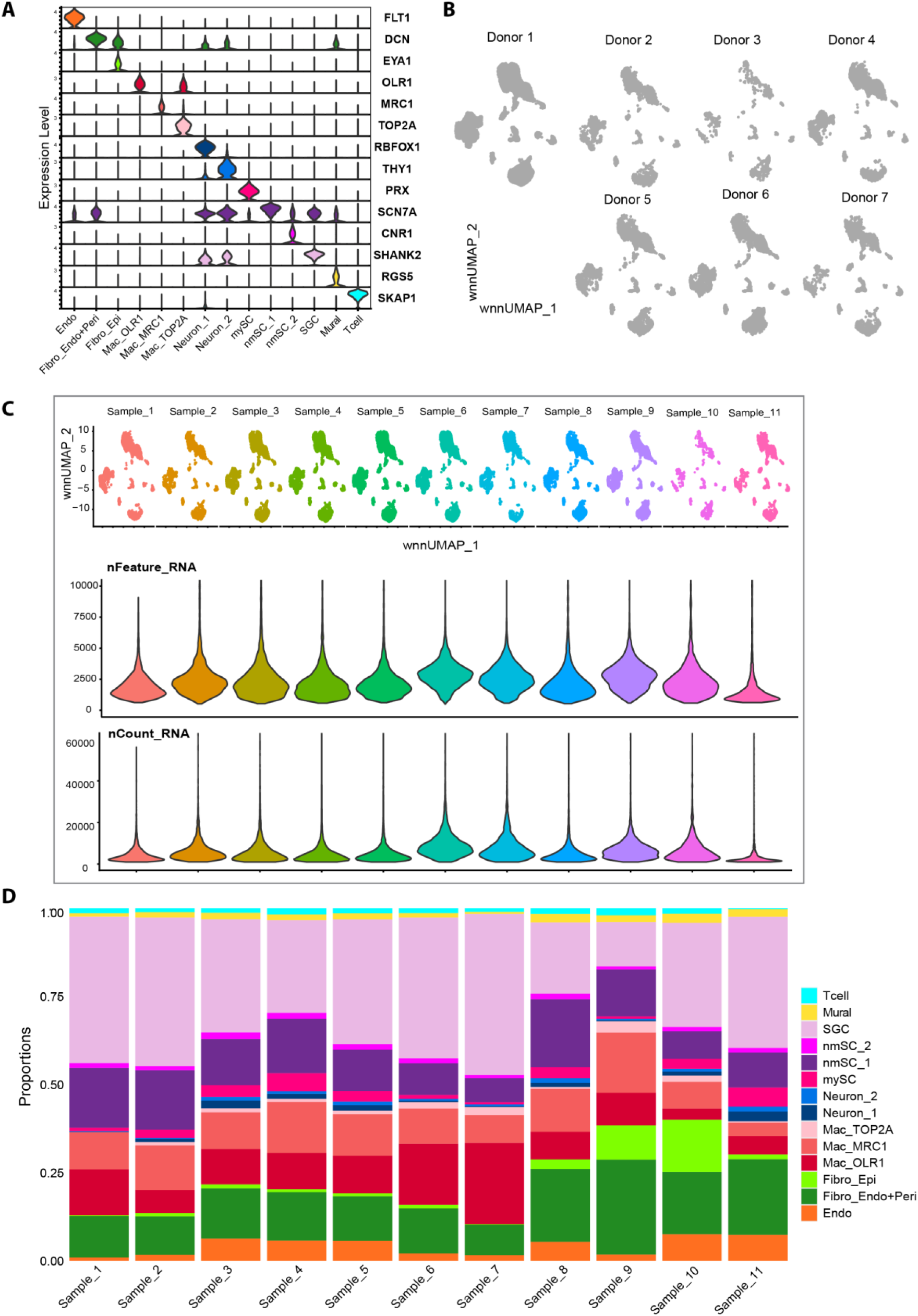
Marker gene expression and cohort information for the 11 hDRG samples. (**A**) Violin plot of marker gene expression levels. (**B**) Cells were visualized using wnnUMAP split by donors. (**C**) Cells were visualized using wnnUMAP split by samples (top) and violin plot of the distribution of total number of features and UMIs for each sample. (**D**) Stacked bar plot of proportion of cells in each sample.

**Figure S3.**
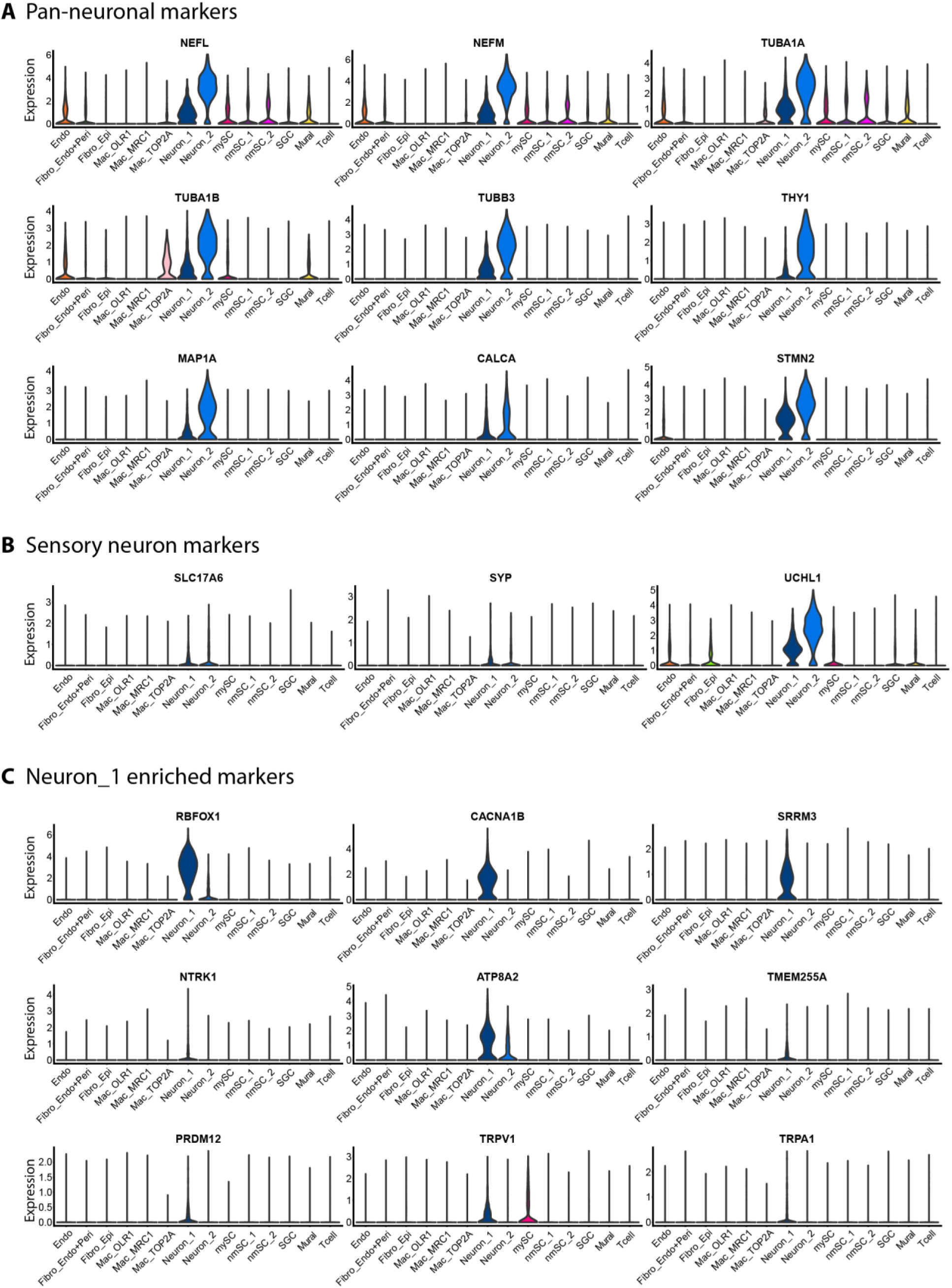
Neuronal marker gene expression. (**A**) Violin plot of pan-neuronal marker gene expression levels. (**B**) Violin plot of sensory neuron marker gene expression levels. (**C**) Violin plot of Neuron_1 marker gene expression levels.

**Figure S4.**
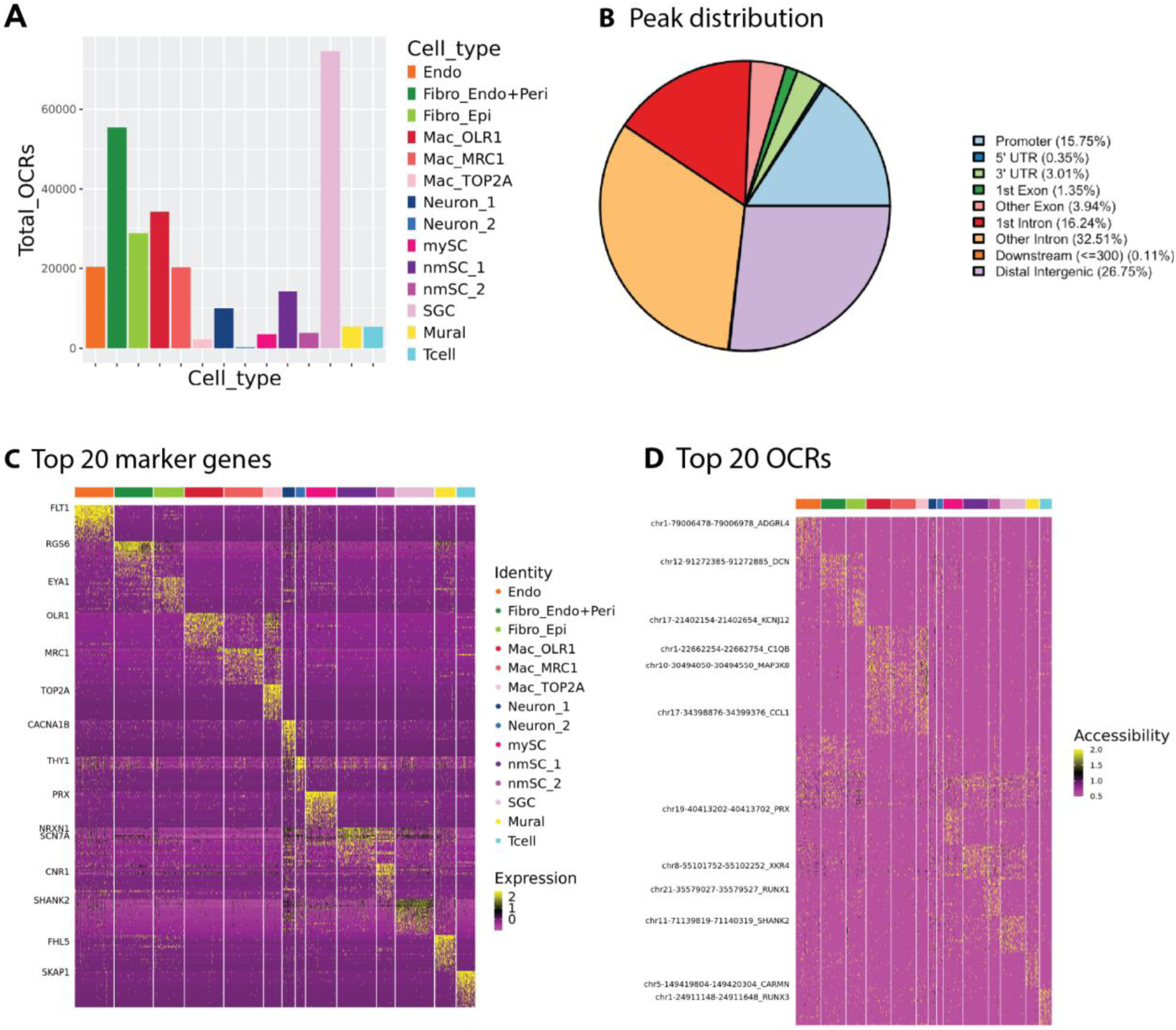
Statistics of open chromatin regions and top marker genes and peaks. (**A**) Bar plot of open chromatin regions (OCRs) identified in each cell type. (**B**) Distribution of OCRs in relation to genomic features. (**C**) Heatmap of top 20 marker genes for each cell type. (**D**) Heatmap of top 20 OCR markers for each cell type.

**Figure S5.**
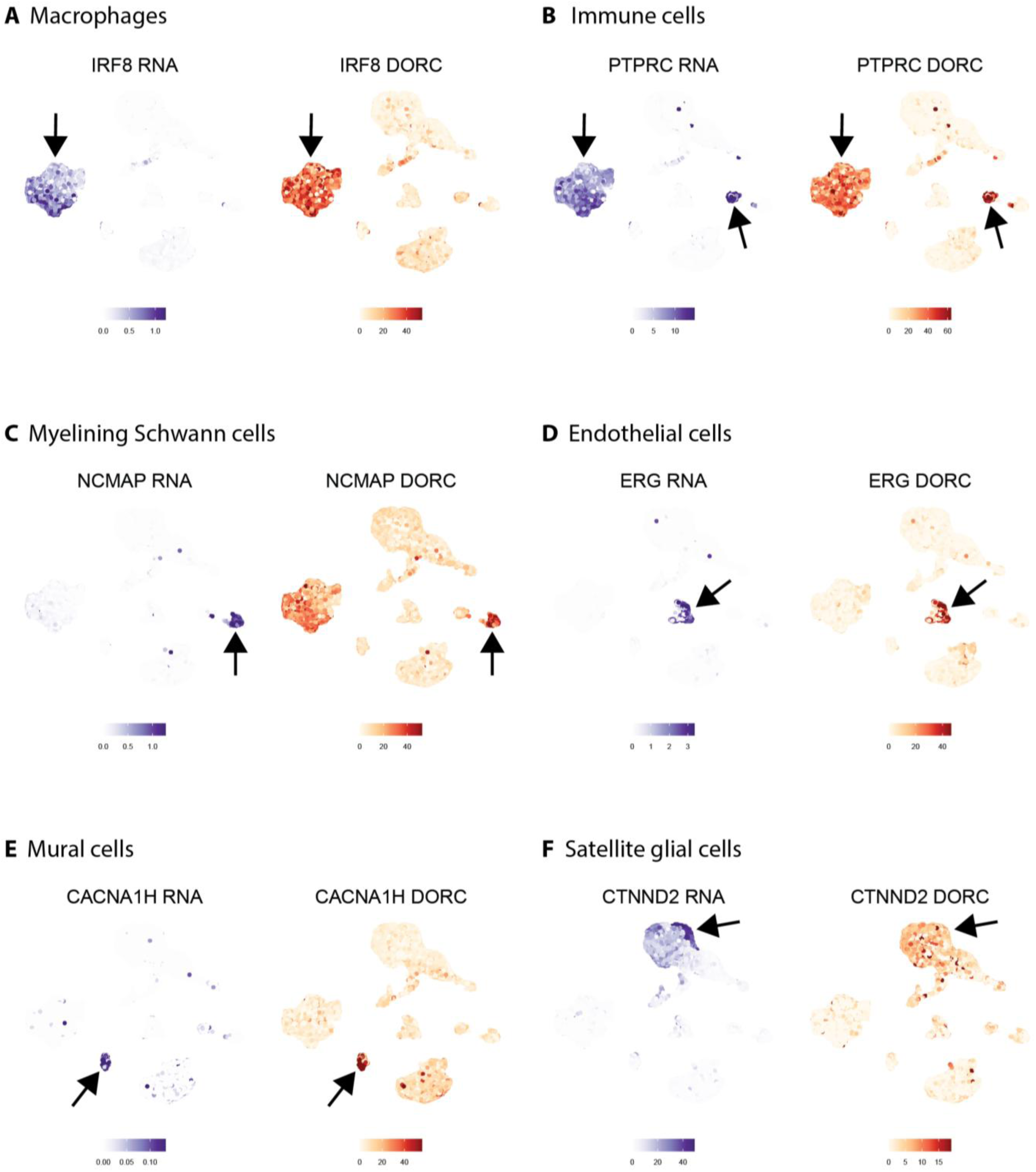
UMAP of DORC accessibility scores (right) and paired RNA expression (left) for marker genes of each cell type.

**Figure S6.**
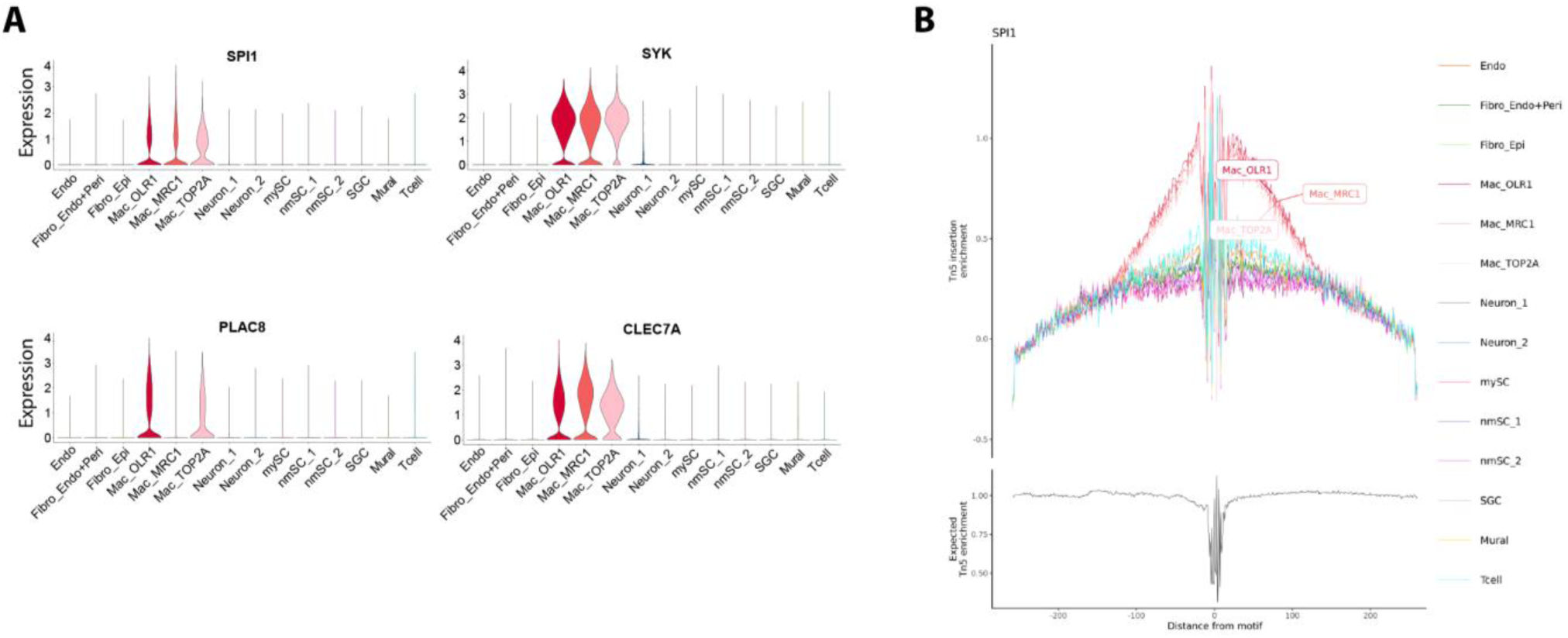
Transcriptional regulatory network analysis for macrophages. (**A**) Violin-plots showing the expression of macrophage-associated DORC genes. (**B**) TF footprinting analysis for SPI1 motifs sites.

**Figure S7.**
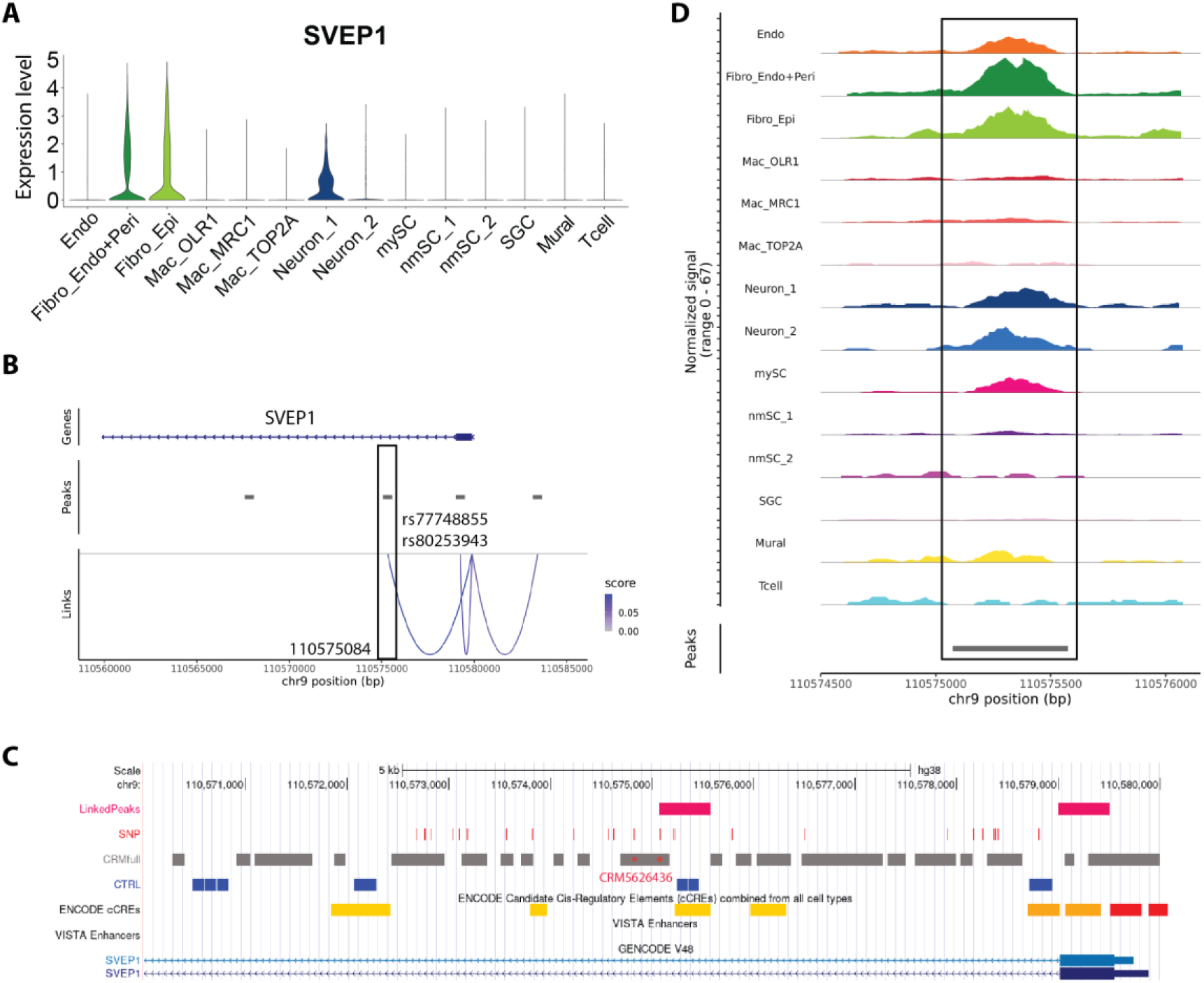
Cell-type specific *SVEP1*-associated risk of pain. (**A**) Violin-plot showing the expression levels of *SVEP1* across all cell populations. (**B**) peak–gene links for *SVEP1* and the location of pain-associated SNPs. (**C**) Genomic view of the snATAC-seq peak significantly associated with *SVEP1*, the location of pain-associated SNPs, VRMOD predicted CRM5626436, and ENCODE cCREs. (**D**) snATAC-seq peak signal.

